# Stress deceleration theory – chronic adolescent stress exposure results in decelerated neurobehavioral maturation

**DOI:** 10.1101/2021.11.21.469381

**Authors:** Kshitij S. Jadhav, Alexander F. Hoffman, Morgane Burgisser, Clara Rosetti, Léa Aeschlimann, Carl R. Lupica, Benjamin Boutrel

## Abstract

Normative development in adolescence indicates that the prefrontal cortex is still under development thereby unable to exert efficient top-down inhibitory control on subcortical regions such as the basolateral amygdala and the nucleus accumbens. This imbalance in the developmental trajectory between cortical and subcortical regions is implicated in expression of the prototypical impulsive, compulsive, reward seeking and risk-taking adolescent behavior. Here we demonstrate that a chronic mild unpredictable stress procedure during adolescence in male Wistar rats arrests the normal behavioral maturation such that they continue to express adolescent-like impulsive, hyperactive, and compulsive behaviors into late adulthood. This arrest in behavioral maturation is associated with the hypoexcitability of prelimbic cortex (PLC) pyramidal neurons and reduced PLC-mediated synaptic glutamatergic control of BLA and nucleus accumbens core (NAcC) neurons that lasts late into adulthood. At the same time stress exposure in adolescence results in the hyperexcitability of the BLA pyramidal neurons sending stronger glutamatergic projections to the NAcC. Chemogenetic reversal of the PLC hypoexcitability decreased compulsivity and improved the expression of goal-directed behavior in rats exposed to stress during adolescence, suggesting a causal role for PLC hypoexcitability in this stress-induced arrested behavioral development.

## Introduction

The limbic circuitry of the mammalian brain is evolutionarily conserved to engender survival (Sokolowski and Corbin, 2012). Key subcortical structures of the limbic circuitry are the basolateral amygdala (BLA), extensively studied for its role in reward processing and assigning value to environmental stimuli (Wassum and Izquierdo, 2015), and the nucleus accumbens, a structure involved in reward and motivation and a primary target of midbrain dopamine projections (Kelley and Berridge, 2002). A later evolutionary addition to this circuit is the prefrontal cortex (PFC), which performs executive functions, such as attention and behavioral flexibility that are critical to problem solving (Yuan and Raz, 2014). The PFC sends substantial inputs to the BLA, and where it exerts top-down control over expression of emotional behavior (Sotres-Bayon and Quirk, 2010), and also to the nucleus accumbens (Phillipson and Griffiths, 1985), where it transmits information critical to goal-directed behavior (Gill et al., 2010).

Mammalian brain development is characterized by protracted periods of excessive synaptogenesis under direct genetic control during the postnatal phase (Petanjek et al., 2008). This excess of synaptic connectivity then undergoes pruning that is influenced by experience, thereby allowing the brain adaptation to the environment (Bick and Nelson, 2016). Importantly, brain development shows a hierarchical pattern with more complex and phylogenetically newer regions such as the PFC maturing later and exhibiting sensitivity to environmental influences into late adolescence (Bourgeois et al., 1994; Huttenlocher et al., 1982). The corticolimbic circuitry also undergoes functional maturation throughout early adolescence into late adulthood (Arruda-Carvalho et al., 2017; Caballero et al., 2014; Cunningham et al., 2002), which is environmentally driven by optimal experiences that promote normal development and facilitating the transition to adulthood (Arnett, 1999). Since subcortical regions mature earlier, compared to the PFC, and the corticolimbic circuitry is still ‘under-construction’ in adolescence, this imbalance is implicated in the expression of quintessential adolescent behaviors such as impulsivity and higher risk taking (Casey et al., 2016; Casey and Jones, 2010). Importantly, experiences during development that are considered outside the realm of “normal”, particularly those involving increased stress, can also influence the pattern of corticolimbic development. Thus, stressful life-experiences, especially during the adolescent period, affect the dendritic structure of neurons in these key brain regions, as well as the functional connectivity among these areas (Leussis et al., 2008; Nooner et al., 2013; Thomason et al., 2015; Wei et al., 2018).

Here, we implement a chronic mild stress procedure during adolescence in male Wistar rats and find that this arrests their behavioral maturation such that they continue to express adolescent-like impulsive, hyperactive, and compulsive behaviors into late adulthood. This behavioral stasis is associated with the hypoexcitability of prelimbic cortex (PLC) pyramidal neurons (PN) and reduced PLC-mediated synaptic glutamatergic control of BLA and nucleus accumbens core (NAcC) neurons that lasts late into adulthood. Chemogenetic reversal of this PLC hypoexcitability decreased compulsivity and improved the expression of goal-directed behavior in rats exposed to stress during adolescence, suggesting a causal role for PLC hypoexcitability in this stress-induced arrested behavioral development.

## Results

Given the central role of the hypothalamo-pituitary-adrenal (HPA) axis in stress regulation (Smith and Vale, 2006), adaptations in corticosterone response to acute stress was assessed in control and CMUS rats. CMUS rats demonstrated lower corticosterone levels on exposure to acute stress, indicating either downregulation or reduced sensitivity to activation of the HPA axis following adolescent chronic stress exposure (Rich and Romero, 2005) (Fig1A)

Aversive experiences may produce neurological changes that can be detected by unconditioned spontaneous behavior. Therefore, we evaluated the effect of adolescent CMUS by testing the performance of these rats in adulthood on the elevated plus maze (EPM) and the open field test (OFT). EPM, tests the behavioral response to the conflict created between a preference for protected areas (closed arms) and the opportunity to explore open environments (open arms). CMUS rats spent significantly more time in the open arms (Control = 23.33 ± 2.74; CMUS = 53.39 ± 2.68, % time on open arm), suggesting a higher level of disinhibited behavior (Fig1B). Moreover, CMUS rats also spent less time in central area of the EPM, suggesting higher risk-taking behavior in the faster decision to explore open arms (Fig1C). The CMUS rats also travelled longer distances in the OFT (Fig1D), indicating elevated motor activity (Control = 36.26 ± 2.62m; CMUS = 57.37 ± 3.84m), which is consistent with increased behavioral disinhibition in CMUS rats since they spent more time (Fig1E) and had more entries in the central region (Fig1F) of the OFT.

Preclinical and human literature suggests that stressful experiences often increase impulsivity (Torregrossa et al., 2012), and adverse childhood experiences are associated with higher impulse control deficits (Barch et al., 2018). Therefore, rats subjected to CMUS in their adolescence were evaluated in the 5-choice serial reaction time task (5CSRTT) in adulthood. In this task, rats must withhold a prepotent motor response to gain access to a reward during a signaled waiting period. While both groups demonstrated similar accuracy (Fig1G), indicating similar learning, the CMUS rats exhibited a lower proportion of omissions (Fig1H), and a significantly higher proportion of premature responses, specifically at the 5s and 7s waiting periods (Control = 23.01 ± 2.72; CMUS = 35.04 ± 3.04; Control = 32.02 ± 4.10; CMUS = 47.64 ± 3.22; % premature responses respectively; Fig1I). A similar accuracy and fewer omissions by the CMUS rats indicated that while the control rats prefer to not respond if they miss the cue light (which potentially reflects a good strategy given the time out as a punishment following an incorrect response), the CMUS rats continue to respond, indicating cognitive inflexibility, and an impulsive phenotype.

The rats were also tested on an attention deficit (AD) task in which they had to maintain attention to a cue light to express an appropriate response. The cue light signaling the correct expected response was decreased in duration over sessions increasing the level of difficulty in detecting the signal. Although both groups exhibited a reduction in the number of correct responses (Fig1J) as the signal duration was decreased, significantly fewer correct responses at the 1s signal duration were observed in the CMUS rats, consistent with impaired attention. Analyzing the incorrect responses in this task (Fig1K) confirmed that adolescent CMUS resulted in lower attention as they consistently display significantly higher incorrect responses at all signal durations (e.g. at 3s signal duration, Control = 7.95 ± 1.26, CMUS = 22.88 ± 4.28; % incorrect responses). Finally, the CMUS rats had a significantly lower proportion of omissions (Fig1L), with this difference being most evident at 0.5s signal duration. Thus, the AD task also demonstrated that, not only do lower correct responses and higher incorrect responses reflect an attention deficit in the CMUS rats, but fewer omissions observed for this group indicated that CMUS rats adopt a higher responding strategy that is non-optimal since it contributes to increased incorrect responses that are punished by a time out period.

The aforementioned behavioral tests investigated motor impulsivity, whereas the delayed discounting task (DDT) measures cognitive impulsivity where we trained rats to choose between a small immediate, or a larger delayed reward with the delay increased within session. Whereas a larger reward was preferred when there was no delay, the choice of larger reward decreased in both the groups as the delay increased (Fig2A). However, the choice of delayed large reward differed between control and CMUS groups, with the latter showing a significant reduction in this choice at a 10s delay (Control = 73.75 ± 6.28; CMUS = 51.67 ± 5.79, % choice for larger reward) (Fig. 2A). This reduced preference for larger reward at the 10 second delay in CMUS rats indicates that they are less able to wait for the larger rewards upon introduction of the delay. However, delays longer than 10s might represent similar difficulty for both groups.

Together, the inability to inhibit a prepotent motor response in the 5CSRTT, the lower proportion of omissions during the AD paradigm, and the inability to choose a larger delayed reward in the DDT, indicate an impulsive phenotype that resulted in inefficient strategies to obtain reward in CMUS rats. However, the motivation to obtain reward was not explicitly assessed in these tasks, and this became the focus of subsequent experiments.

In a test of motivation for reward, the CMUS rats exhibited a significantly higher preference for saccharine and much lower variability on this measure than controls (Control = 82.25 ± 7.14, CMUS = 98.20 ± 0.19, % saccharine preference) (Fig2B). CMUS rats also consumed higher volumes of saccharine, compared to water (Control = 31.95 ± 10.64; CMUS = 67.68 ± 6.43, saccharine to water ratio) (Fig2C). The increased saccharine preference was also associated with an increased motivation for reward, as CMUS rats had significantly higher break points in the progressive ratio task (Control = 79.29 ± 12.48; CMUS = 132.8 ± 17.78, lever presses until rats stop responding) (Fig2D). The CMUS rats were also more persistent at reward taking when lever presses for saccharine reward were followed by mild foot shocks (Control = 11.44 ± 4.91; CMUS = 87.67 ± 33.94, lever presses) (Fig2E) indicating increased compulsive reward taking in the CMUS rats.

Finally, compulsivity is also defined by the inability to adapt responding to the changing value of reward, reflecting a shift toward habitual, rather than goal-directed behavior. Therefore, we used a behavioral task to assess responding to two distinct rewards, one that was devalued and the other non-devalued, followed by extinction. Rats demonstrating normal goal directed behavior focus their responding on a lever previously associated with the non-devalued reward (valued lever) and ignore the lever associated with the devalued reward (Balleine and Dickinson, 1998). After the devaluation procedure, control rats showed preference for the valued lever, whereas CMUS rats showed similar a preference for the valued and devalued lever [Control (Valued: 49.14 ± 4.74, Devalued: 30.08 ± 4.74); CMUS (Valued: 35.80 ± 4.56, Devalued: 53.68 ± 14.75), % baseline lever presses/min], indicating reduced goal-directed behavior and increased habitual responding (Fig2F).

Having identified these behavioral changes caused by stress exposure during adolescence, we next sought to identify changes in neuronal function that may underlie these effects. For these electrophysiological experiments, rats included in the CMUS group underwent the stress experience as explained in the methods, followed by evaluation in the EPM. Like our previous experiment, the CMUS rats spent significantly more time in the open arms of the EPM, compared the controls (Suppl Fig1A).

As stressful experiences during adolescence are known to impair PFC development (Andersen and Teicher, 2008; 2009; Arnett, 1999; Baker et al., 2013; Eiland et al., 2012; Spear, 2000), we examined the post-synaptic properties of neurons in key corticolimbic regions immediately after the stress procedure at PND50 using whole-cell patch electrophysiology (Fig3A). In layer 5 (L5) PNs of the PLC there was no effect of CMUS on the resting membrane potential (Control = −65.74 ± 1.09 mV, CMUS = −64.96 ± 0.85 mV). However, the mean input resistance of L5 PLC PNs was significantly smaller (Fig3B), and the smallest depolarizing current to elicit an action potential (rheobase) was significantly higher in CMUS rats (Control = 555.2 ± 56.22pA, CMUS = 808.2 ± 51.92pA) (Fig3C). PLC neurons from CMUS rats also showed a significantly lower frequency of action potential (AP) discharge across a range of currents injected through patch electrodes [e.g., 250pA current AP frequency, Control = 39.54 ± 2.39Hz, CMUS = 24.07 ± 1.83Hz] (Fig3D 1-2). Further, when the contribution of small conductance calcium-activated potassium channel (Sk) function to after-hyperpolarization (AHP) amplitudes was measured, we found no difference in the sAHP amplitude (Control = 0.56 ± 0.14mV, CMUS = 0.84 ± 0.16mV), and a significant increase in the mAHP amplitude (Control = 4.02 ± 0.25mV, CMUS = 5.05 ± 0.35mV) in the CMUS rats PLC PNs (Suppl Fig1B 4-5). This increase in mAHP could act to decrease AP discharge frequency, thereby contributing to increased inhibition of these cells. We also found that the frequency of spontaneous synaptic glutamate EPSCs (sEPSCs) was lower in neurons from CMUS rats (Control = 2.98 ± 0.41Hz, CMUS = 1.77 ± 0.35Hz) (Suppl Fig1B 2,3), suggesting a decrease in excitatory synaptic drive of these cells. Together, these data identify several cellular adaptations that underlie L5 PLC neuron hypoexcitability in rats exposed to the adolescent stress procedure.

Given the central influence of the BLA on the adolescent corticolimbic circuitry (Walker et al., 2017), the effects of stress exposure during adolescence on BLA PNs was investigated (Fig3E). In contrast to the hypoexcitability of PLC PNs after stress, BLA PNs showed increased excitability in CMUS rats. Thus, while there was no difference in the resting membrane potential (Control = −63.53 ± 0.5mV, CMUS = −63.19 ± 0.61mV), the mean rheobase was significantly smaller (Control = 116.8 ± 14.34pA, CMUS = 50.44 ± 4.96pA), and input resistance and AP discharge with current injection (350pA current step AP frequency, Control = 26.16 ± 1.55Hz, CMUS = 35.48 ± 1.96Hz) were significantly increased (Fig. 3F, Fig. 3G). Also, both mAHP (Control = 4.1 ± 0.32mV, CMUS = 2.51 ± 0.26mV) and sAHP (Control = 0.52 ± 0.09mV, CMUS = −0.03 ± 0.16mV) were significantly decreased after CMUS. We also found that mean sEPSC frequency was higher in BLA PNs from the CMUS rats (Control = 2.59 ± 0.37 Hz, CMUS = 5.06 ± 0.66 Hz) (Suppl Fig1C 4,5, Suppl Fig1C 2,3). Thus, in stark contrast to PLC neurons, exposure to stress in adolescence increased the excitability of BLA neurons.

Finally, since CMUS rats exhibited enhanced reward sensitivity in our behavioral tasks, and the NAcC is involved in motivation and receives glutamatergic projections from both the BLA and PLC, we investigated the electrophysiological properties of medium spiny neurons (MSN) in this brain region (Fig3I). We found that there were no differences in the properties of NAcC MSNs in between the CMUS and control groups. Thus, input resistance (Fig3J), rheobase (Fig3K), AP frequency, Sk channel function (Suppl Fig1D 4,5) and sEPSC frequency and amplitude were all unchanged by CMUS (Suppl Fig1D 2,3).

To determine the duration of the effects of stress exposure in adolescence, additional cohorts of rats underwent the CMUS procedures and the neuronal properties were evaluated at PND90. All post-synaptic differences observed between CMUS and control rats at PND50 were observed at PND90, indicating that these changes are not transient but long-term and potentially irreversible. (Suppl Fig2).

Having established that the stressful life experiences in adolescence alters the excitability of the PLC and BLA PNs and that NAcC MSNs are unchanged, we asked whether communication among these brain regions was affected by stress. To do this, we used optogenetics (Suppl Fig3) combined with *invitro* electrophysiology to examine changes in excitatory synaptic pathways among these regions. There was a tendency towards smaller optically elicited EPSCs (oEPSCs) following CMUS (Fig4A). Thus, the amplitudes of oEPSCs in both groups increased with increased 473nm light intensity (input-output relationship, I-O) but at different rates and optogenetic stimulation activated significantly weaker oEPSCs in CMUS rat BLA PNs at 10 mW intensity (Control = 400.9 ± 56.56 pA, CMUS = 234.1 ± 38.66 pA) (Fig 4B,C). Additionally, a significantly smaller proportion of BLA PNs from the CMUS rats fired APs driven by light-activated EPSPs (oEPSPs) from PLC axons (Control = 63.63%; CMUS = 27.27%, proportion of neurons firing AP) (Fig4D). The strength of glutamatergic PLC inputs to NAcC MSNs (Fig4E) was significantly lower in the CMUS rats as indicated by the shift in the I-O curve [oEPSC amplitude at 10mW light intensity (Control = 559.9 ± 91.96pA, CMUS = 253.1 ± 44.16pA)] (Fig4F,G). Also, a significantly smaller proportion of NAcC MSNs fired an APs in response to light-activated glutamatergic PLC afferents in the CMUS rats (Control = 80%; CMUS = 37.5%, neurons firing AP) (Fig4H). These data indicate that stress exposure during adolescence reduces the strength of glutamatergic projections from the PLC to both BLA and NAcC, and results in a significant reduction in synaptically-driven AP firing probability. This likely indicates that information transfer from the PLC to both BLA and NAcC is significantly reduced in the aftermath of adolescent stress.

Further investigation of the corticolimbic circuitry demonstrated that the strength of BLA glutamate inputs to PLC L5 PNs (Fig4I) was lower in the CMUS rats, as indicated by the smaller slope of the oEPSC I-O curve [oEPSC amplitude at 10mW light intensity (Control = 225.8 ± 36.47pA, CMUS = 98.23 ± 11.66pA)] (Fig 4J,K). This decreased excitatory synaptic drive of PLC PNs also resulted in a significantly reduced probability of these cells firing APs in response to synaptic BLA input in the CMUS rats (Control = 43.75%; CMUS = 7.14%, neurons firing AP) (Fig4L). This suggests that information transmission to the PLC from the BLA is reduced by exposure to stress in adolescence and this could impair PFC maturation that is dependent on this input (Tottenham and Gabard-Durnam, 2017). Finally, I-O curves also showed that the strength of BLA glutamate inputs to the NAcC MSN were significantly increased in CMUS rats (Control = 168.7 ± 35.59pA, CMUS = 342.2 ± 50.04pA)] (Fig. 4M,N,O). Moreover, this augmented excitatory input dramatically increased the probability that the BLA-driven oEPSPs would evoke AP discharge in NAcC MSNs (Control = 7.69%; CMUS = 100%, neurons firing AP) (Fig4P). Thus, the enhanced strength of glutamatergic projections from the BLA to NAcC following adolescent stress indicates stronger control of the NAcC by the BLA in the CMUS rats. This, coupled with the changes in BLA and PLC PN postsynaptic properties (Fig3), indicates that NAcC MSNs are rendered more responsive to BLA inputs, compared to those from the PLC.

Having investigated the strength of the synaptic connectivity between the three key regions of the corticolimbic circuitry, we next investigated the effects of adolescent stress on short- and long-term synaptic plasticity. All of the glutamatergic projections demonstrated similar short-term plasticity, as evaluated using paired pulse activation of oEPSCs (Fig5E-H). Therefore, we next evaluated long-term spike-timing dependent plasticity (STDP) in which synaptic glutamate release is paired with firing of the postsynaptic neuron. In BLA PNs, the STDP protocol resulted in long-term depression (LTD) of PLC to BLA glutamatergic oEPSCs in both CMUS and control groups. However, the magnitude of LTD was significantly larger in the CMUS group when measured 30 min after the STDP protocol was initiated (Two way repeated measures ANOVA [TW-RM-ANOVA], group effect, F(1,22)= 5.96, p=0.023) (Fig5I, Fig5M). Similarly, spike-timing-dependent LTD of the glutamatergic projection from PLC to NAcC MSNs was significantly larger in the CMUS group 30 min after initiating the STDP protocol Control = 72.61 ± 6.14 pA, CMUS = 53.46 ± 4.25pA) (Fig5J, Fig5N). Further, there was stronger LTD of the glutamatergic projection from the BLA to PLC PN in the CMUS group (TW-RM-ANOVA, group effect, F(1,21)= 8.36, p=0.0087) over the last 5 minutes of the experiment (Control = 81.01 ± 14.25pA, CMUS = 46.34 ± 4.5pA) (Fig5K-O). Finally, LTD in the projections from BLA to NAcC was not affected by adolescent stress (Fig5L, 5P). These data suggest that CMUS is associated with an increased susceptibility to long-term depression of PLC to BLA, BLA to PLC, and PLC to NAcC projections, but not in the BLA to NAcC projection.

The electrophysiology experiments demonstrated that stress exposure during adolescence results in the hypoexcitability of PLC PNs, combined with weaker top-down control from the PLC to the BLA and NAcC. To determine whether this weakened PLC output is involved in the altered behaviors observed after adolescent stress, we used chemogenetics to reverse the PLC hypoexcitability during behavioral tasks. Prior to surgical infusion of the AAV8-hM3D-Gq in the PLC, we confirmed the disinhibited behavior on the EPM, hyperactivity on the OFT, and motor impulsivity on the 5CSRTT in the CMUS rats (Suppl Fig4A-E). Following the CMUS procedure, rats were assigned to one of 4 groups (i) AAV-hM3D-Gq in PLC + vehicle (ii) AAV-hm3D-Gq in PLC + clozapine, (iii) Sham surgery in PLC + vehicle, (iv) sham surgery in PLC + clozapine (Suppl Fig4F-J). After training on a FR1 schedule for saccharine, rats from each group were injected with 0.1mg/kg clozapine, or an equivalent volume of vehicle, 30 minutes before the compulsivity testing and active lever presses for 0.2% saccharine during foot shock were measured. CMUS rats infused with AAV8-hM3D-Gq in the PLC and injected with clozapine had fewer active lever presses during foot shock, compared to CMUS rats receiving inactive sham injection or those receiving virus injection in PLC and vehicle injection (active lever presses per group (i) 79 ± 22.82, (ii) 19.75 ± 3.54, (iii) 95.57 ± 30.02, (iv) 137.85 ± 33.35) (Fig 6C). Moreover, the number of active lever presses was not affected by virus infusion or clozapine injection in CMUS rats (Suppl Fig4K), suggesting no changes in general motor behavior. Together the data suggest that the PLC is recruited during conflictual situations and that stress during adolescence suppresses PLC PN excitability resulting in ineffectual compulsive behavior.

Finally, these rats were tested in a goal-directed devaluation task using a Latin square design in which they each received vehicle or clozapine injections in different sessions. Thus, each rat underwent 5-minutes of extinction after 1hr of sensory specific devaluation and was entered into the next phase of the study. They then received 0.1mg/kg of clozapine, or in a separate session, an equivalent volume vehicle injection. A two-way repeated measures ANOVA, with clozapine injection and devaluation effect as two factors, was run separately for the control rats (n=7), CMUS inactive sham rats (n=10) and CMUS Gq virus injected rats (n=11). Three control rats, 4 CMUS inactive virus surgery rats and 5 CMUS Gq virus injected rats were deleted from analysis due to either low baseline lever pressing behavior or biased consumption test (Suppl data: goal.xlsx). Controls rats demonstrated appropriate goal directed behavior, irrespective of receiving either clozapine (Valued: 49.01 ± 12.68; Devalued: 23.38 ± 7.37, % of baseline lever press/min) or vehicle (Valued: 46.15 ± 6.96; Devalued: 28.32 ± 7.51, % of baseline lever press/min) (Fig6D1). However, CMUS rats who underwent PLC inactive virus surgery chose the devalued and valued reward levers to the same extent, indicating impaired goal-directed behavior, irrespective of clozapine (Valued: 69.81 ± 14.34; Devalued: 66.24 ± 13.69, % of baseline lever press/min) or vehicle injection (Valued: 61.38 ±11.07; Devalued: 59.81 ± 17.21, % of baseline lever press/min) (Fig6D2). However, CMUS rats with AAV-hM3D-Gq in PLC and clozapine injection showed appropriate goal directed behavior (valued: 46.89 ± 5.74; devalued: 24.96 ± 3.52, % of baseline lever press/min), compared to that measured with vehicle injection (Valued: 50.60 ± 5.21; Devalued: 51.45 ± 9.79, % of baseline lever press/min) (Fig6D3). These data suggest that stress exposure in adolescence renders PLC PNs hypoexcitable and this leads to aberrant goal directed behavior that can be reversed by chemogenetically increasing the activity of these neurons. Therefore, the data indicate that PLC neuron hypofunction is likely causal in the behavioral deficits caused by adolescent stress.

## Discussion

The theory of developmental programming posits that environmental influences during sensitive periods of development can influence the structure and function of physiological systems, and potentially contribute to persistent neural changes (Padmanabhan et al., 2016). Thus, the consequences of stressful experiences beyond the boundaries of normal expectancy can be magnified during a sensitive period like adolescence because it represents a time of critical neural development marked by high cellular plasticity (Spear, 2000) and maturation of brain circuits (Giedd, 2004).

It is well-established in both clinical (Cauffman et al., 2010; Figner et al., 2009) and preclinical studies (Andrzejewski et al., 2011) (Adriani and Laviola, 2003) that adolescents exhibit impulsive behavior. Moreover, preclinical studies show that stressful adolescent experiences are associated with impulsive behaviors even in adulthood (Baarendse et al., 2013; Torregrossa *et al*., 2012), and this is confirmed by clinical observations (Barch *et al*., 2018; Schalinski et al., 2018). Studies in rodents also show that chronic stress results in decreased attention in adulthood (Novick et al., 2013), and that perturbation of social context results in a disinhibited behavior in adulthood (Shao et al., 2009; Watt et al., 2009). In addition to these consequences of stress, adolescents are known to engage in risky behavior characterized by sensation seeking (Doremus-Fitzwater and Spear, 2016), and in the absence of intervening challenge this behavior declines during maturation (Romer et al., 2017). The increased impulsive behavior in adulthood following adolescent stress suggests that this experience might slows the normal neurobehavioral maturational process. Therefore, we suggest that the high impulsivity, hyperactivity, disinhibited behavior, and increased reward seeking we report in rats undergoing CMUS in our study is a consequence of this altered developmental process. More formally, we hypothesize that stress experienced during adolescence slows down the behavioral maturation process resulting in expression of behavior reminiscent of adolescence in adulthood (Figure 7).

This ‘behavioral syndrome’ consists of higher emotionality, diminished forethought and impaired executive control, all characteristics previously associated with PFC and BLA function. In our study, stress during adolescence weakened glutamatergic projections to PLC PNs and decreased intrinsic excitability of neurons in this structure. This likely contributes to impaired information integration in these neurons and reduced propagation of this activity to subcortical regions. However, we also found that adolescent CMUS had an opposite effect on BLA PNs. Thus, BLA cells received stronger glutamatergic input, and exhibited higher intrinsic excitability, potentially contributing to increased propagation of information to downstream brain regions, such as the NAcC (Figure 7). This opposing pattern of adolescent stress effects on the PLC and BLN in CMUS rats is similar that reported in humans, where amygdala activity is increased and PFC activity decreased during emotional regulation in adults that experienced peri-adolescent stress (Kim et al., 2013; McLaughlin et al., 2015).

In contrast to the effects of adolescent stress on BLA and PLC neurons, we observed no changes in properties of NAcC MSNs. This may be due to MSNs reaching functional maturity before the CMUS procedure. It has been suggested that the NAcC acts to integrate information coming from the PFC and BLA, with the BLA transmitting emotionally salient information (Stuber et al., 2011) and the PFC input promoting executive control over inappropriate action (Kalivas et al., 2005). In our study the glutamatergic projection from PLC to NAcC was weakened, and that from BLA to NAcC strengthened following adolescent stress, suggesting that this loss of cortical control would shift more toward BLA-mediated emotional control of the NAcC output. Human brain imaging research also shows decreased functional coupling between PFC and ventral striatum during risky decision making in adolescent humans (Telzer et al., 2015), as well as stronger resting state connectivity between amygdala and accumbens in adolescents that have experienced trauma (Nooner *et al*., 2013). In general, our findings agree with these changes and provide putative mechanisms for changes observed in these imaging studies. We propose based on these findings that stress-induced corticolimbic imbalance tips the scale towards a greater influence of BLA glutamatergic projections on NAcC MSNs, potentially increasing the emotional bias of information impinging on the NAcC. This, together with the weakened PFC driven inhibitory control, likely contributes to the hyperactive, disinhibited, impulsive phenotype observed in our rats. Further, as the PFC is necessary for goal-directed behavior expression (Buschman and Miller, 2014), and decreased top-down control likely affects this function, perhaps not surprisingly we also demonstrate that goal-directed behavior is reduced in the CMUS rats.

It has recently been shown that model-based control, which is a reflection of goal directed behavioral control, improves with age during adolescence and this is associated with higher functional coupling between the PFC and ventral striatum in humans (Vaghi et al., 2020). However, adolescents who do not exhibit normative maturation of PFC-ventral striatum connectivity exhibit higher compulsivity (Vaghi *et al*., 2020). In our study, stress exposure in adolescence resulted in impaired goal-directed behavior and increased compulsive behavior that was associated with reduced PLC-NAcC connectivity and reduced glutamatergic synaptic control. Thus, the deceleration of corticolimbic circuit development after adolescent stress resulted in the expression of adolescent like behavior in adulthood in our CMUS rats. Other evidence supporting this idea comes from the finding that the trajectory of development of goal-directed behavioral control is steeper in early human adolescence (Vaghi *et al*., 2020), which is consistent with the developmental stage at which our CMUS paradigm was implemented in rats (PND21). Thus, our results support the idea that there is a critical developmental window when exposure to stress has a large impact upon adult behavior.

The idea that stress decreases top-down control of behavior is well supported in our investigation where it is demonstrated that there is weaker glutamatergic control of BLA PNs that originates from the PLC. As the maturation of this neural circuitry is associated with increased connectivity during the transition from adolescence to adulthood (Arruda-Carvalho *et al*., 2017; Hare et al., 2008), and exposure to trauma in adolescent humans results in lower PFC-amygdala functional connectivity (Thomason *et al*., 2015), we suggest that our findings support the stress-induced deceleration of corticolimbic circuitry proposed in this model (Figure 7).

While it is often stated that the PFC is modulated by the environment, at the neural level, one of the major sources of influence over the PFC is the amygdala, which takes in environmental information and transmits it to the PFC, thus shaping its development especially during adolescence (Tottenham and Gabard-Durnam, 2017). This is supported by previous literature which shows that the connectivity between BLA, and PFC continues to develop from adolescence to adulthood (Caballero *et al*., 2014; Cunningham *et al*., 2002; Nooner *et al*., 2013). Thus, the decreased strength of synaptic glutamatergic connections between BLA and PFC observed in our study supports the proposal that stress during adolescence also decreases the functional maturation of this circuit (Figure 7).

Finally, our study also provides causal evidence that the hypoexcitability of PLC PNs caused by adolescent stress is involved in the expression of the compulsive and diminished goal-directed behavior. Thus, when the hypoactive PLC was chemogenetically activated, these behavioral changes were normalized. Our control experiments also confirm the specificity of this manipulation because clozapine alone did not affect appropriate baseline lever press behavior, indicating that the drug did not alter motor behavior. These data also show that the imposition of a conflictual state, such as that seen with the presence of shock during reward seeking in the compulsivity test, or in a value assessment seen after devaluation in goal-directed behavioral test, is necessary for the recruitment of the PLC and to observe these behavioral deficits. Thus, the hypoexcitability the PLC following adolescent stress was causally associated with this behavioral expression since chemogenetic activation of this brain structure increased behavioral control as shown by reduced compulsivity and improved goal-directed behavior.

## Conclusion

Stress is ubiquitous in life and short-term neurocircuit adaptations are essential for adjustment to temporary stressors. Whereas these adaptations can be beneficial in dealing with short-term stressors, they can be detrimental to developing brain circuits and behavior when the stress is sustained over a longer period of time. Here, we demonstrate that chronic and sustained mild stressful experiences in adolescence can result in the development of neurobehavioral adaptations that could increase the long-term risk of detrimental behaviors and psychiatric disorders. Therefore, understanding the neural mechanisms contributing to circuit adaptations resulting in the development of this hyperactive, impulsive, compulsive phenotype is important to developing treatments and interventions. Future experiments should focus on potential preventive and curative strategies to counter the deleterious effects of adolescent stress.

---

***<u>Figure 1: Adolescent exposure to stress attenuated corticosterone response to acute stress and induced a disinhibited, hyperactive, impulsive and inattentive phenotype</u>***

*Data is presented as mean ± SEM, number of rats mentioned after statistical analysis [Control=C, CMUS=S]*.

**(A)** Stress exposure during adolescence attenuated the corticosterone response to acute stress even if the basal corticosterone level as well as time to return to baseline was similar in both the group. *[Two way repeated measures ANOVA (TW-RM-ANOVA), group effect (F(1,22) = 5.64, p=0.02), time effect (F(2,44) = 81.52, p<0.0001) and interaction effect (group X time) (F(2,44) = 5.26, p=0.0089), post hoc Sidak’s test (t12= 3.95, p=0.0006). **C:12,S:12]***

**(B)** CMUS rats also showed decreased time spent on the open arm of the EPM *[unpaired T test (t55 = 7.823, p<0.0001, **C:29, S:28]*** reflecting their disinhibited behaviour **(C)** with lower time spent in the central zone of the EPM alluding to their higher risk taking behavior *[unpaired T test, t55 = 2.035, p=0.045, **C:29, S:28]*** **(D)** CMUS rats also exhibited hyperactivity indicated by longer distance travelled in OFT *[Unpaired T test, t29= 4.47, p=0.0001, **C: 15,S: 16.]*** **(E)** with more time spent in the central zone *[Unpaired T test, t29= 3.34, p=0.0023), **C: 15,S: 16.]*** **(F)** and higher entries in the central zone of the OFT both indicating disinhibited behavior *[Unpaired T test, t29= 3.9, p=0.0005), **C: 15,S: 16]*** On the 5CSRTT, stress exposure during adolescence did not affect the learning ability indicated by **(G)** similar accuracy in both the groups *[TW-RM-ANOVA, no group effect (F(1,31) = 0.004, p=0.94), no group X duration of waiting, (F(2,62) = 1.224, p=0.3), **C: 15,S: 18]***, **(H)** The increased impulsivity was indirectly indicated by the lower proportion of omissions in the 5CSRTT *[TW-RM-ANOVA, no group effect (F(1,31) = 0.74, p=0.39), significant effect of duration of waiting (F(2,62) = 22.44, p<0.0001), significant interaction effect (F(2,62) = 5.46, p=0.0065), significant post hoc Sidak’s test at 5 seconds duration of waiting (t93 = 2.868, p=0.015), C: 15,S: 18.]* **(I)** which was then confirmed by higher proportion of premature responses *[TW-RM-ANOVA, significant group effect F(1,31) = 8.46, p=0.0066, significant duration of waiting effect (F(2,62) = 22.94, p<0.0001) significant post hoc Sidak’s test, 5s (t93 = 2.33, p=0.04) and 7s (t93 = 3.02, p=0.0095), **C: 15,S: 18.]***

Finally, the decreased attention in the CMUS rats was observed in the **(J)** lower proportion of correct responses on the AD test *[TW-RM-ANOVA: significant group effect (F(1,30) = 4.25, p=0.04), significant interaction effect (group X signal duration, F(3,90) = 3.40, p=0.021), significant signal duration effect (F(3,90) = 459.1, p<0.0001), post-hoc Sidak’s test (t120 = 3.457, p=0.003), **C: 16,S: 16.]*** along with **(K)** higher proportion of incorrect responses as well *[TW-RM-ANOVA, group effect (F(1,30) = 16.9, p=0.0003), signal duration effect (F(3,90) = 145, p<0.0001), post hoc Sidak’s test (3s, [t120= 3.55, p=0.0022], 1s [t120=4.19, p=0.0002], 0.5s [t120= 3.68, p=0.0014], 0.2s [t120= 2.72, p=0.029]), **C: 16,S: 16.]*** The CMUS rats also demonstrated **(L)** lower omissions on the AD test reflective of their impulsive behavior *[TW-RM-ANOVA, group effect (F(1,30) = 6.07, p=0.0197), singal duration effect (F(3,90) = 15.06, p<0.0001)), (post hoc Sidak’s test (t120 = 3.483, p=0.0028)), **C: 16,S: 16.]***

***<u>Figure 2: Stress exposure in adolescence also increased impulsive choice along with aberrant attraction for sweetened reward in CMUS rats</u>*.**

**(A)** Introduction of a delay on the Delay Discounting task resulted in CMUS rats to shift more towards choosing the smaller reward compared to the control rats reflecting higher cognitive impulsivity *[(TW-RM-ANOVA Group effect (F(1,10) = 0.97, p=0.34), Delay effect (F(4,40) = 52.36, p<0.0001)), (group X duration of delay, (F(4,40) = 4.25, p=0.0058)), (post hoc Sidak’s test, (t50 = 2.75, p=0.04)). **C: 6, S: 6**.]*

In the two bottle choice test the CMUS rats showed an aberrant attraction for the sweetened reward as reflected in **(B)** higher Saccharine preference *[(Mann-Whitney test, U=9, p=0.0021), **C: 9, S: 10.]*** and **(C)** higher saccharine to water ratio *[Unpaired T test, t12=2.54, p=0.0258), **C: 9, S: 10]***

**(D)** CMUS rats also demonstrated higher motivation for saccharine in progressive ratio task *[Mann-Whitney test, U=9, p=0.0117, **C: 8, S: 6.]***

(**E)** Introduction of a conflictual situation by delivery of a shock after lever press for saccharine showed higher compulsive behavior in CMUS rat *[Mann-Whitney U test, U=5, p=0.0117), **C:8, S: 6]***

**(F)** Finally, the CMUS rats demonstrated lower goal directed behavior indicating inability to adapt responding based on the value of a reward *[TW-RM-ANOVA, no group effect (F(1,12) = 0.32, p=0.57), no devaluation effect (F(1,12) = 0.005, p=0.94) but a significant interaction effect group X devaluation (F(1,12) = 5.54, p = 0.036). (Simple effect analysis, t6=5.174, p=0.0021), (Simple effect analysis, t6 = 1.173, p = 0.28), **C: 6, S: 7.]***

***<u>Figure 3: Effects of adolescent stress exposure on the Postsynaptic electrophysiological properties of L5 PLC PN, BLA PN and NAcC MSN at PND50</u>*.**

*Data is presented as mean ± SEM, number of cells (number of rats) mentioned after statistical analysis [Control=C, CMUS=S]*.

**(A)** Pictorial representation of the location of the investigated PLC L5 PN. Stress exposure during adolescence **(B)** decreased the input resistance of the L5 PLC PN of the CMUS rats *[Unpaired t test, t60= 4.701, p=0.0001, **C: 31(5), S: 31(5)]***. **(C)** A higher injected current was needed to elicit an AP (higher rheobase) *[Unpaired t test, t53= 3.31, p=0.0017, **C: 27(5), S: 28(5)]*** along with **(D1)** lesser number of elicited AP frequency *[(TW-RM-ANOVA, group effect, (F(1,53) = 29.69, p<0.0001), current effect, (F(5,265) = 391.7, p<0.0001), group X injected current, (F(5,265) = 21.98, p<0.0001)) observed at all injected current strength (post hoc Sidak’s test 50pA[p=0.033], 100, 150, 200 and 250pA p<0.0001 for all), **C: 27(5), S: 28(5)]*** both reflect a reduced excitability of L5 PLC PN. **(D2)** Representative traces for D1 *(Control in blue and CMUS in orange*)

**(E)** Pictorial representation of the location of the investigated BLA PN. Stress exposure during adolescence **(F)** increased input resistance in BLA PN of CMUS *rats [Unpaired t test, t71= 4.981, p<0.0001, **C: 39(5), S: 34(5)]*** **(G)** A lower rheobase *[Unpaired T test, t66= 4.37, p<0.0001, **C: 34(5), S: 34(5)]*** and **(H1)** higher number of elicited AP frequency reflected an increased excitability of BLA PN *[(TW-RM-ANOVA, group effect (F(1,54) = 26.07, p<0.0001), current effect (F(7,378) = 505.0, p<0.0001, group X injected current, (F(7,378) = 13.41, p<0.0001)) and higher number of AP were fired in CMUS rat neurons at all injected current (post hoc Sidak’s test, (50, 100, 150, 200, 250, 300 and 350pA, p<0.0001 for all)), **C: 27(5), S: 29(5)]*** **(H2)** Representative traces for H1 *(Control in blue and CMUS in orange*)

**(I)** Pictorial representation of the location of the investigated NAcC MSN. Stress exposure in adolescence did not affect the **(J)** input resistance *[Mann-Whitney U test, U= 64, p=0.67, **C: 10(5), S: 16(5)]***, **(K)** rheobase *[Unpaired T test, t22= 0.029, p=0.97, **C: 10(5), S: 16(5)]*** or ***(L1)*** the excitability of the NAcC MSN *[(TW-RM-ANOVA, no group effect (F(1,22) = 0.098, p=0.75), current effect (F(5,110) = 128.9, p<0.0001), no group X injected current (F(5,110) = 0.6027, p=0.69)), **C: 10(5), S: 16(5)]***. **(L2)** Representative traces for L1 *(Control in blue and CMUS in orange*)

***<u>Figure 4: Effects of stress exposure during adolescence on the synaptic connectivity within the corticolimbic circuitry</u>***

*Data is presented as mean ± SEM (except panels D,H,L,P which are represented as absolute number of neurons), number of cells (number of rats) mentioned after statistical analysis [Control=C, CMUS=S]*.

**(A)** Pictorial representation of the site of injection of AAV5-CaMKII-ChR2-eYFP in the PLC and the site of recorded PN in the BLA with optically elicited glutamate release. **(B)** Representative traces of the oEPSCs for (A). **(C)** Stress exposure during adolescence resulted in a weaker glutamatergic projections from PLC PN to BLA PN *[TW-RM-ANOVA, group effect, (F(1,27) = 4.02, p=0.055)), (light intensity effect, (F(8,216) = 54.59, p<0.0001)), (group X light intensity effect, (F(8,216) = 2.99, p=0.0034), Post hoc Sidak’s test, p=0.038 at 10mW, **C: 11(4), S: 11(5)] (D)*** Further, lower proportion of BLA PN fired an AP due to weaker oEPSP elicited by the glutamate release from the PLC PN terminals *[Fisher’s exact test, (p=0.04), **C: 11(4), S: 14(5)]***

**(E)** Pictorial representation of the site of injection of AAV5-CaMKII-ChR2-eYFP in the PLC and the site of recorded NAcC MSN with optically elicited glutamate release **(F)** Representative traces of the oEPSCs for (E). **(G)** Stress exposure during adolescence resulted in a weaker glutamatergic projections from PLC PN to NAcC MSN *[TW-RM-ANOVA, group effect, (F(1,36) = 13.40, p=0.0008), light intensity effect, (F(8,288) = 35.85, p<0.0001), group X light intensity effect, (F(8,288) = 1.685, p=0.1017). The oEPSC amplitude was significantly lower in the CMUS rats’ NAcC MSN at light intensity of 3, 4, 5, 6, 7, 8, 9 and 10mW (post-hoc Sidak’s test, p<0.01 for all). **C: 15(5), S: 23(6)] (H)*** Further, lower proportion of NAcC MSN fired an AP due to weaker oEPSP elicited by the glutamate release from the PLC PN terminals *[Fisher’s exact test, (p=0.0029)). **C: 15(5), S: 16(6)]***

**(I)** Pictorial representation of the site of injection of AAV5-CaMKII-ChR2-eYFP in the BLA and the site of recorded PN in the PLC with optically elicited glutamate release **(J)** Representative traces of the oEPSCs for (I). **(K)** Stress exposure during adolescence resulted in a weaker glutamatergic projections from BLA PN to PLC PN *[(TW-RM-ANOVA, group effect, (F(1,31) = 10.42, p=0.0029)), (light intensity effect, (F(8,248) = 32.99, p<0.0001)), (group X light intensity effect, (F(8,248) = 4.503, p<0.0001)). The oEPSC amplitude were smaller in the CMUS group PLC layer 5 PN at light intensities 4, 5, 6, 7, 8, 9 and 10 mW (Sidak’s test, p<0.05 for all). **C: 14(4), S: 17(6)] (L)*** Further, lower proportion of PLC PN fired an AP due to weaker oEPSP elicited by the glutamate release from the BLA PN terminals *[(Fisher’s exact test, (p=0.039)). **C: 16(5), S: 14(6)]***

**(M)** Pictorial representation of the site of injection of AAV5-CaMKII-ChR2-eYFP in the BLA and the site of recorded NAcC MSN with optically elicited glutamate release. **(N)** Representative traces of the oEPSCs for (M). **(O)** Stress exposure during adolescence resulted in a stronger glutamatergic projections from BLA PN to NAcC MSN *[TW-RM-ANOVA, group effect, (F(1,36) = 6.203, p=0.0017)), (light intensity effect, (F(8,288) = 27.92, p<0.0001)), (group X light intensity effect, (F(8,288) = 5.195, p<0.0001)), The oEPSC amplitude were larger in the CMUS group NAcC MSN at light intensity of 7, 8, 9, 10mW (Sidak’s test, p<0.05 for all). **C: 15(5), S: 23(5)] (P)*** Further, higher proportion of NAcC MSN fired an AP due to larger oEPSP elicited by the glutamate release from the BLA PN terminals *(Fisher’s exact test, (p<0.0001)). **C: 14(5), S: 15(5)]***

***<u>Figure 5: Short term and Long term synaptic plasticity in the corticolimbic circuitry post adolescent stress exposure</u>***

*Data is presented as mean ± SEM. Number of cells (number of rats) mentioned after statistical analysis [Control=C, CMUS=S]*.

**(A-D)** Pictorial representations of the site of injection of AAV5-CaMKII-ChR2-eYFP and the site of recorded neurons with optically elicited glutamate release

**(E-H)** All the investigated corticolimbic glutamatergic projections demonstrated comparable Paired pulse ratio in both the groups (representative traces in inset).[(**E) PLC → BLA***: TW-RM-ANOVA, no group effect (F(1,20) = 0.47, p=0.49) or interaction effect (F(4,80) = 0.91, p=0.45) and only an inter-stimulus interval (ISI) effect (F(4,80) = 35.65, p<0.0001). **C: 11(4), S: 10(5),*** **(F) PLC → NAcC**: *TW-RM-ANOVA, no group effect (F(1,31) = 0.208, p=0.65) or interaction effect (F(4,124) = 0.51, p=0.72) and only an ISI effect (F(4,124) = 43.18, p<0.0001). **C: 15(4), S: 18(6),*** **(G) BLA → PLC:** *TW-RM-ANOVA, no group effect (F(1,25) = 0.92, p=0.34) or interaction effect (F(4,100) = 0.35, p=0.84) and only an ISI effect (F(4,100) = 23.67, p<0.0001). **C: 15(4), S: 12(6),*** **(H) BLA → NAcC:** *TW-RM-ANOVA, no group effect (F(1,34) = 2.55, p=0.11) or interaction effect (F(4,136) = 0.19, p=0.93) and only an ISI effect (F(4,136) = 90.09, p<0.0001)). **C: 15(5), S: 21(5)**]*

***(I-L)*** The long term synaptic plasticity within the investigated corticolimbic circuitry was studied using the Spike time dependent protocol (STDP).The projection from **(I)** PLC→ BLA showed a stronger LTD in the CMUS group *[TW-RM-ANOVA time effect (F(59,1298)=4.46, p<0.0001), group effect(F(1,22)=5.964, p=0.023,no interaction effect (F(59,1298)=1.107, p=0.27). **C: 11(4), S: 13(4)]***, while the **(J)** PLC → NAcC projection showed similar LTD *[TW-RM-ANOVA time effect (F(59,1239)=5.28, p<0.0001), no group effect (F(1,21)=2.873, p=0.1), no group X time effect (F(59,1239)=0.93, p=0.61). **C: 11(4), S: 12(5)]***. **(K)** Further BLA → PLC glutamatergic projection demonstrated a stronger LTD in the CMUS rats *[TW-RM-ANOVA time effect (F(59,1239)=4.33, p<0.0001), group effect (F(1,21)=8.36, p=0.0087)*) with steeper fall in oEPSC amplitude in the CMUS group neurons *(group X time effect (F(59,1239)=1.572, p=0.0043). **C: 10(4), S: 13(5)]*** while **(L)** the LTD in the BLA → NAcC projection was similar in both the groups *[TW-RM-ANOVA, time effect (F(59,1221)=8.33, p<0.0001)), no group differences (no group effect(F(1,19)=1.41, p=0.24, no group X time effect (F(59,1121)=0.63, p=0.98). **C: 10(4), S: 11(5)]***.

**(M-P)** LTD for the investigated corticolimbic projections was also measured as a fall in the average oEPSC amplitude in the last 5 minutes of STDP when expressed as a percentage of baseline oEPSC amplitude. While **(M)** the PLC → BLA *[Unpaired T test, t22=2.52, p=0.01. **C: 11(4), S: 13(4)]***, **(N)** PLC→ NAcC *[Unpaired T test, t21=2.59, p=0.01. **C: 11(4), S: 12(5)]*** and BLA → PLC *[Unpaired T test, t21=2.58, p=0.01, **C: 10(4), S: 13(5)]*** glutamatergic projection showed stronger LTD in the CMUS rats, the BLA → NAcC glutamatergic projection demonstrated similar LTD *[Unpaired T test, t19=0.68, p=0.5. **C: 10(4), S: 11(5)]***.

***<u>Figure 6: Chemogenetic activation of PLC PN rescues effects of adolescent stress exposure on compulsive and goal directed behavior</u>***

*Data is presented as mean ± SEM, number of rats mentioned after statistical analysis*

**(A)** Pictorial representation of site of injection of the AAV9-hM3D-Gq in the PLC

**(B)** Representative immunofluorescence image showing the expression of AAV8 excitatory DREADD (pAAV-CaMKIIa-hM3D(Gq)-mCherry) injected in the PLC (right hemisphere)

**(C)** Rats injected with PN activating DREADDs in the PLC decreased their lever pressing behavior compared to rats undergoing Sham surgery on chemogenetic activation *[Two way ANOVA, (F(1,26)=7.662, p=0.01))],* while there was no difference between rats injected with clozapine or vehicle *[Two way ANOVA, (F(1,26)=0.12, p=0.73)]*. However, the two factors interacted indicating an influence of clozapine injection depending on the presence or absence of PLC PN activating DREADD (DREADD viral injection in PLC X clozapine injection) *[F(1,26)=4.355, p=0.04)]*. The CMUS rats injected with PN activating DREADD in PLC decreased their lever pressing behaviour when administered clozapine as compared to rats undergoing Sham surgery were administered clozapine *[Post-hoc Sidak’s test, t=3.43,df=26, p=0.004]*. Further, the Sham surgery rats performed similarly irrespective of being injected with either clozapine or vehicle *[Simple effect analysis (t=0.9398, df=12), p=0.36)]*. Importantly, PLC PN activation by clozapine in CMUS rats decreased the lever pressing behavior of rats in the compulsivity paradigm *[Simple effect analysis (p=0.0002**)].[CMUS PLC Gq DREADD–vehicle (n=8), CMUS PLC Gq***

***<u>Supplementary Figure 1:</u>***

*Data is presented as mean ± SEM, number of cells (number of rats) mentioned after statistical analysis [Control=C, CMUS=S]. Experiments were conducted at PND50. Representative traces are presented in inset*

**(A)** CMUS rats exhibited disinhibited behavior on the EPM as they spent significantly more time on the open arms of the EPM as compared to the controls confirming our previous experiments *[Unpaired T test, t86= 7.2, p<0.0001). **C=46, S=42]***

**(B1)** Pictorial representation of the location of the investigated PLC L5 PN. **(B2)** There was a decreased frequency of sEPSC impinging on L5 PN of the PLC in CMUS rats which was significantly lower as compared to the control group *[Unpaired T test, t41=2.199, p=0.033). **C: 21(5), S: 22(5)] (B3)*** However, there was no difference in the amplitude of the sEPSC impinging on these neurons *[Unpaired t test, t41= 0.98, p=0.32, **C: 21(5), S: 22(5)]*** indicating that it is mainly a presynaptic effect with potentially less number of glutamatergic projections on to these pyramidal neurons. **(B4)** The SK channel function measured by mAHP was enhanced in the L5 PN of the PLC of the CMUS rats compared to the control rats *[Unpaired t test, t47= 2.32, p=0.024, **C: 24(5), S: 25(5)]*** which could account for decreased excitability in CMUS rats PN. **(B5)** However, there was no difference in the sAHP in the L5 PN of the PLC in both the groups *[Unpaired t test, t47= 1.24, p=0.21, **C: 24(5), S: 25(5)]***

**(C1)** Pictorial representation of the location of the investigated BLA PN. **(C2)** There was a increased frequency of sEPSC impinging on PN of the BLA in CMUS rats which was significantly higher as compared to the control group *[Unpaired T test, t46=2.91, p=0.0055, **C: 20(5), S: 28(5)] (C3)*** However, there was no difference in the amplitude of the sEPSC impinging on these neurons *[Unpaired T test, t46= 0.86, p=0.39, **C: 20(5), S: 28(5)]*** indicating that it is mainly a presynaptic effect with potentially higher number of glutamatergic projections on to these pyramidal neurons. **(C4)** The SK channel function measured by mAHP *[Unpaired t test, t47= 2.32, p=0.024, **C: 24(5), S: 25(5)]*** and **(C5)** sAHP *[Unpaired T test, t40= 2.79, p=0.008, **C: 19(5), S: 23(5)]*** was decreased in the PN of the BLA of the CMUS rats compared to the control rats which could account for increased excitability in CMUS rats PN.

**(D1)** Pictorial representation of the location of the investigated NAcC MSN. There was no difference in either the **(D2)** frequency *[Unpaired T test, t24= 0.56, p=0.46, **C: 10(5), S: 16(6)]*** or the **(D3)** amplitude *[Unpaired T test, t24= 0.49, p=0.55, **C: 10(5), S: 16(6)]*** of the sEPSC impinging on the MSN of the NAcC of CMUS and control rats. Further, SK channel function as measured by **(D4)** mAHP *[Unpaired T test, t24= 0.48, p=0.65, **C: 10(5), S: 16(6)]*** and **(D5)** sAHP was similar of NAcC MSN in both the groups *[Unpaired T test, t24= 0.35, p=0.92, **C**: **10(5), S: 16(6)]***.

*Supplementary Figure 2:*

*Data is presented as mean ± SEM, number of cells (number of rats) mentioned after statistical analysis [Control=C, CMUS=S]. Experiments were conducted at PND90*.

**(A)** Pictorial representation of the location of the investigated PLC L5 PN. The PLC L5 PN in the CMUS rats had **(B)** lower input resistance compared to the control rats *[Unpaired t test, t33= 3.52, p=0.0013, **C: 20(4), S: 15(4)**]*. This coupled with **(C)** higher rheobase (least current needed to elicit an AP) pointed towards reduced excitability of L5 PLC PN [*Unpaired t test, t28= 3.73, p=0.0008, **C: 18(4), S: 12(4)**]*. **(D)** The lower number of AP fired as the injected current was increased confirmed the reduced excitability of L5 PLC PN *[TW-RM-ANOVA, group effect, (F(1,25) = 9.79, p=0.0044), current effect, (F(5,125) = 103.1, p<0.0001), group X injected current, (F(5,125) = 60.61, p<0.0001) observed at all injected current strength (post hoc Sidak’s test 50pA[p=0.033], 150, 200 and 250pA p<0.01 for all), **C: 17(4), S: 11(4)**]*. One possible reason for this reduced excitability could be due to the enhanced SK channel function measured by higher **(E)** mAHP *[Unpaired t test, t24= 3.11, p=0.0048, **C: 14(4), S: 12(4)**]* and **(F)** sAHP *[Unpaired t test, t24= 2.95, p=0.0069, **C: 14(4), S: 12(4)**]* amplitudes in the CMUS rats L5 PLC PN as compared to the controls

**(G)** Pictorial representation of the location of the investigated BLA PN. The BLA PN in the CMUS rats had **(H)** higher input resistance compared to the control rats *[Unpaired t test, t28= 2.88, p=0.0074, **C: 18(6), S: 12(4)**]*. This coupled with **(I)** lower rheobase (least current needed to elicit an AP) pointed towards increased excitability of BLA PN *[Unpaired T test, t27= 2.87, p=0.0077, **C: 17(6), S: 12(4)**]*. **(J)** The higher number of AP fired as the injected current was increased confirmed the increased excitability of L5 PLC PN *[TW-RM-ANOVA, group effect (F(1,27) = 13.40, p=0.0011), current effect (F(7,189) = 462.5, p<0.0001)), group X injected current, (F(7,189) = 8.53, p<0.0001) and higher number of AP were fired in CMUS rat neurons at all injected current (post hoc Sidak’s test, (100, 150, 200, 250, 300 and 350pA, p<0.01 for all)), **C: 17(6), S: 12(4)**]*. One possible reason for this higher excitability could be due to the reduced SK channel function measured by **(K)** lower mAHP amplitude in the BLA PN of the CMUS rats[*Unpaired T test, t22= 4.905, p<0.0001) which could account for increased excitability in CMUS rats BLA PN. **C: 12(6), S: 12(4**)]*. **(L)** The sAHP amplitude was however similar in the BLA PN of both the groups *[Unpaired T test, t22= 1.96, p=0.061, **C: 12(6), S: 12(4)***]

**(M)** Pictorial representation of the location of the investigated NAcC MSN. The **(N)** input resistance *[Unpaired T test, t42= 0.04, p=0.68, **C: 23(8), S: 21(8)**]*, **(O)** rheobase *[Unpaired T test, t24= 1.19, p=0.24), **C: 14(8), S: 12(8)**]* as well as the **(P)** excitability *[TW-RM-ANOVA, no group effect (F(1,23) = 2.47, p=0.12), current effect (F(4,92) = 137.5, p<0.0001), no group X injected current interaction effect (F(4,92) = 1.96, p=0.16), **C: 13(8), S: 12(8)**]* of the MSN of the NAcC of CMUS and Control rats were similar. Predictably, there was no difference in the **(Q)** mAHP *[Unpaired T test, t24= 0.1.28, p=0.20, **C: 14(8), S: 12(8)**]* and **(R)** sAHP *[Unpaired T test, t24= 0.34, p=0.73, **C: 14(8), S: 12(8)**]* amplitude in the MSN of NAcC in CMUS and control rats

**Supplementary Figure 3:**

**(A)** Representative image of PLC infusion of AAV5-CaMKII-ChR2-eYFP

**(B)** Representative image of BLA infusion of AAV5-CaMKII-ChR2-eYFP

**Supplementary Figure 4:**

Data is presented as mean ± SEM, number of rats mentioned after statistical analysis [Control=C, CMUS=S, CMUS sham vehicle =SSV, CMUS sham clozapine = SSC, CMUS Gq Vehicle = SGV, CMUS Gq clozapine = SGC]. Results explained in terms of CMUS rats

**(A) CMUS rats demonstrated the classical higher disinhibited behavior** as expressed by the higher amount of time spent on the open arm in the EPM paradigm as compared to the control rats (Unpaired T test, t38=5.038, p<0.0001). **C=10, S=30**

**(B) CMUS rats demonstrated hyperactivity** as demonstrated by longer distances travelled by them as compared to the control rats (Mann Whitney U test, U=71, p=0.012). **C=10, S=30**

**(C) CMUS rats showed a similar proportion of correct responses on the 5CSRTT** as compared to control rats (TW-RM-ANOVA, group effect, F(1,38)=3.27, p=0.078), as well as no interaction effect (F(2,76)=1.98, p=0.144). While there was an effect of time of delay (F(1.887, 71.72)=20.18, p<0.0001), post hoc analysis showed that there proportion of correct responses were significantly lower in the CMUS group only at time of delay of 10 seconds (t37.98=3.127, p=0.01). This shows that the both the groups on an average learnt the paradigm similarly. **C=10, S=30**

**(D) CMUS rats showed a similar proportion of omission responses on the 5CSRTT** as compared to control rats (TW-RM-ANOVA group effect, F(1,38)=3.218, p=0.08), as well as no interaction effect (F(2,76)=2.36, p=0.1012). While there was an effect of time of delay (F(1.901, 72.25)=6.14, p<0.0001) indicating that both the groups continued to learn and hence showed lower proportion of omission responses as training sessions increased, post hoc analysis showed that the proportion of omission responses were similar in both the groups at all time points. **C=10, S=30**

**(E) CMUS rats showed a higher proportion of premature responses on the 5CSRTT** as compared to control rats (TW-RM-ANOVA, group effect, F(1,38)=8.432, p=0.0061) indicating higher motor impulsivity in the CMUS as previously demonstrated by us. While there was an effect of time of delay (F(1.872, 71.12)=39.75, p<0.0001) indicating that both the groups showed higher number of premature responses as the time of delay increased. There was however no interaction effect (F(2,76)=0.11, p=0.89). Post hoc Sidak’s test showed that proportion of premature responses were significantly higher at 5s time of delay (t11.84 = 3.02, p=0.031). **C=10, S=30**

**(F) All four CMUS groups showed similar behavior on EPM.** To determine baseline similarity between rats subsequently split for the compulsivity paradigm, we ran a two way ANOVA for only the CMUS rats with future PLC Gq injection as one factor and future clozapine injection as another factor. This showed that all the four groups were similar to each other for the EPM paradigm at baseline. (Gq injection factor, F(1,26)=1.293, p=0.265) (clozapine injection factor, F(1,26)=0.0028, p=957). **SSV= 7, SSC= 7, SGV= 8, SGC= 8.**

**(G) All four CMUS groups undergoing surgery showed similar behavior on the OFT.** To determine baseline similarity between rats subsequently split for the compulsivity paradigm, we ran a two way ANOVA for only the CMUS rats with future PLC Gq injection as one factor and future clozapine injection as another factor. This showed that there was no difference between Gq and Sham surgery rats (Gq injection factor, F(1,26)=1.48, p=0.23). While there was a clozapine factor effect (F(1,26)= 8.115, p=0.008), there was no interaction effect (F(1,26)=0.3096, p=0.58). Further, post hoc analysis showed that there was no difference between all the four groups except vehicle injected CMUS Gq rats and clozapine injected CMUS sham rats (t26=4.06, p=0.037). So, on a whole all the four groups were more or less similar to each other for their OFT measured hyperactivity. **SSV= 7, SSC= 7, SGV= 8, SGC= 8.**

**(H) All four CMUS groups undergoing PLC viral infusion showed similar accuracy on the 5CSRTT.** To determine baseline similarity between rats subsequently split for the compulsivity paradigm, we ran a three way ANOVA for only the CMUS rats with PLC Gq injection as one factor, future clozapine as the second factor and the time of delay the third factor. While there was an effect of time of delay (F(2,52)=9.755, p=0.0003), there was no effect of Gq injection (F(1,26)=0.013, p=0.9095), no effect of clozapine factor (F(1,26)=0.002, p=0.96), no interaction effect of time of delay X Gq injection (F(2,52)=0.83, p=0.44), no interaction effect of time of delay X clozapine factors (F(2,52)=0.73, p=0.48), marginal interaction effect of Gq injection X clozapine factors (F(1,26)=4.63, p=0.041) and finally no interaction effect of time of delay X Gq injection X Clozapine factors (F(2,52)=1.37, p=0.26). A post hoc Sidak’s multiple comparison test showed there was no significant difference in the proportion of correct responses of the four groups at the same time of delay indicating that all the four groups learnt the paradigm similarly. **SSV= 7, SSC= 7, SGV= 8, SGC= 8.**

**(I) All four CMUS groups undergoing PLC viral infusion showed similar omissions on the 5CSRTT**. To determine baseline similarity between rats subsequently split for the compulsivity paradigm, we ran a three-way ANOVA for only the CMUS rats with PLC Gq injection as one factor, future clozapine as the second factor and the time of delay the third factor. While there was an effect of time of delay (F(1.732,45.04)=4.522, p=0.02), an effect of Gq injection (F(1,26)=4.77, p=0.03), an effect of clozapine factor (F(1,26)=5.08, p=0.03), no interaction effect of time of delay X Gq injection (F(2,52)=0.04, p=0.95), no interaction effect of time of delay X clozapine factors (F(2,52)=2.23, p=0.11), no interaction effect of Gq injection X clozapine factors (F(1,26)=0.02, p=0.87) and finally no interaction effect of time of delay X Gq injection X Clozapine factors (F(2,52)=0.18, p=0.83). A post hoc Sidak’s multiple comparison test showed there was no significant difference in the proportion of omission responses of the four groups at the same time of delay indicating that all the four groups learnt the paradigm similarly. **SSV= 7, SSC= 7, SGV= 8, SGC= 8.**

**(J) All four CMUS groups undergoing PLC viral infusion showed similar premature responses on the 5CSRTT**. To determine baseline similarity between rats subsequently split for the compulsivity paradigm, we ran a three-way ANOVA for only the CMUS rats with PLC Gq injection as one factor, future clozapine as the second factor and the time of delay the third factor. While there was an effect of time of delay (F(1.919,49.90)=47.36, p<0.0001), no effect of Gq injection (F(1,26)=1.89, p=0.18), no effect of clozapine factor (F(1,26)=1.44, p=0.24), no interaction effect of time of delay X Gq injection (F(2,52)=0.88, p=0.41), no interaction effect of time of delay X clozapine factors (F(2,52)=3.007, p=0.058), no interaction effect of Gq injection X clozapine factors (F(1,26)=1.65, p=0.21) and finally no interaction effect of time of delay X Gq injection X Clozapine factors (F(2,52)=0.36, p=0.69). A post hoc Sidak’s multiple comparison test showed there was no significant difference in the proportion of premature responses of the four groups at the same time of delay indicating that all the four groups showed similar baseline motor impulsivity. **SSV= 7, SSC= 7, SGV= 8, SGC= 8**

**(K) No effect of clozapine on motor behavior in CMUS rats.** To determine no effect of clozapine on motor behavior, CMUS rats that underwent PLC viral infusion or sham surgery were tested in a Latin square design during FR1 training for 0.2% saccharine with either 0.1 mg/kg clozapine or equivalent volume of vehicle injection. A two way repeated measures ANOVA showed that there was no effect on either PLC surgery or clozapine injection on lever pressing behavior during baseline training. (No effect of Gq injection (F(1,28)=0.47, p=0.49), no effect of clozapine factor (F(1,28)=0.11, p=0.73), no interaction effect of PLC surgery X clozapine injection (F(1,28)=0.24, p=0.87). CMUS sham= 14, CMUS Gq = 16.

## Methodology

### Animals

Male Wistar rats were bred in-house at the Center for Psychiatric Neuroscience animal facility (breeders ordered from Charles River, France). They were at PND21 and weighed 50-60 g at the beginning of the experiment and they were randomly assigned to the CMUS or the control group. All experiments were performed in accordance with the Swiss Federal Act on Animal Protection and the Swiss Animal Ordinance and were approved by the cantonal veterinary office (authorization 3047 to Benjamin Boutrel).

All electrophysiological experiments were approved by the Institutional Care and Use Committee of the National Institute on Drug Abuse Intramural Research Program (NIDA-IRP), National Institutes of Health (NIH), and conducted in accordance with the Guide for the Care and Use of Laboratory Animals provided by the NIH and adopted by the NIDA-IRP. The Male Wistar pups for these experiments were ordered from Charles-River Lab when they were 14 days old along with a dam (6 pups per dam). After one-week (PND21) (acclimatization period), pups were weaned, and group housed (3 per cage). Then they were randomly assigned to the CMUS or the control group.

In both settings the rats were kept in reversed 12-h light/dark cycle (lights off at 0830 hours) and housed in controlled temperature and humidity conditions.

### Chronic Mild Unpredictable Stress (CMUS) procedure

Rats assigned to the CMUS groups were subjected to the CMUS procedure as mentioned in Table 1.

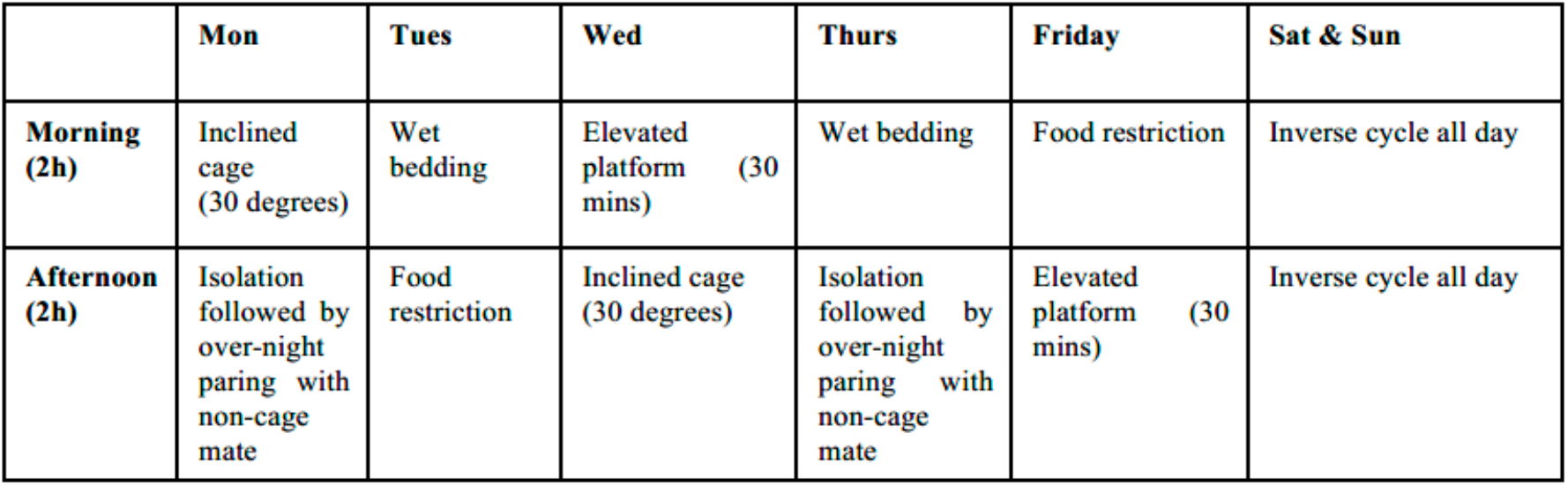

Control rats were handled regularly but not subjected to the CMUS procedure.

### Corticosterone response to acute stress

The baseline blood sample was collected from the tail vein of the rats 1 day prior to the acute stress paradigm at 1200 hours. Next day, the rats were subjected to the elevated platform (12 × 12 × 100 cm above the ground) at 1200 hours for a period of 30 mins followed immediately by tail vein blood collection. The rats were then returned to their home cage and the final tail vein blood sample was collected 1 hour after the end of the acute stress procedure. The blood samples (200–300 μl) were systematically collected using heparin Microvette CB 300 (Sarstedt AG, Sevelen, Switzerland) after tail vein incision. Blood samples were then centrifuged at 4°C for 20 minutes at 4500 rpm and frozen at −20°C until analysis. Plasma samples were analyzed using a commercial enzymatic-immuno assay kit (Corticosterone EIA kit, Enzo Life Sciences, Lausen, Switzerland) following the manufacturer instructions.

### Elevated plus maze

The EPM apparatus was positioned 50 cm above the floor and divided in four arms: two “closed arms” enclosed by plastic wall (500×100×425 mm), and two “open arms” without walls. In the center, a small open arena (100×100 mm) allowed access to each arm. Luminosity was fixed at 20 lux in the open arms, and 10 lux in the closed arms. Rats were placed at the center of the arena facing one of the open arms and their exploratory activity was recorded for 5 min. The percentage of time spent in the open arms was used as the index of disinhibition, and locomotor activity was further assessed. The video tracking system AnyMaze 6.0 (Stoelting Europe, Dublin, Ireland) was used to track the activity of the rats.

### Open Field Test

Three open field arenas were used for this procedure. Each arena was a cylinder made of grey Plexiglas with a diameter of 70cm and a height of 30cm. The three cylinders were put on a large grey Plexiglas table. The luminosity was approximately 24 lux at the center of the arena. The three arenas were recorded simultaneously by a vertically mounted video camera linked to a computer in the same room. The video tracking system AnyMaze 6.0 (Stoelting Europe, Dublin, Ireland) was used to track the activity of the rats. Three animals from the same cage were placed each in a different arena against the wall. The animal freely explored the arena for 30 minutes. At the end of the experiment the rats were put back in their home cage and the arenas were cleaned with Deconex and dried before bringing in the next set of animals.

### 5-Choice serial reaction time task (5CSRTT): Impulsive action

The 5CSRTT is used to measure motor impulsivity or impulsive action in rats. Six operant chambers (30.5 × 24.1 × 29.2 cm) were used. Each chamber was individually enclosed in a wooden cubicle, which was equipped with an exhaust fan for ventilation as well as served as a white noise speaker for sound attenuation. Each operant chamber has a steel grid and a tray with bedding. On the left side of the chamber, five holes (25×25 mm) for nosepokes were present horizontally and separated by 25mm from each other. Opposite these five holess, on the right side of the chamber, was the food receptacle, located 20mm above the grid. Lights for the nosepokes, food receptacle and chamber ceiling were present. Sucrose pellets (Dustless precision pellet 45mg, rodent purified diet, Bioserv, Frenchtown, NJ) were used during the entire experiment as a reward delivered upon each correct response. The sucrose pellets were stored in a receptacle outside the cage and for each correct response, one sucrose pellet was released to the food receptacle.

Rats were maintained at 85% of their age matched body weight and were fed 1 hour after the end of the session each day. The 5CSRTT was carried out following five training steps.

#### Training 1

Rats underwent magazine training and learned to perform a nosepoke in the food receptacle to receive one sucrose pellet as a reward. The rats were considered to have reached the training criteria once they reached 50 pellets in 30 mins.

#### Training 2

Next, when a nosepoke in the food receptacle occurred, a light stimulus appeared randomly in one of the five holes and stayed lit until another nosepoke in this hole occurred, even if a nosepoke in any of the four other unlit holes was emitted. Once a nosepoke in the lit hole occurred, this light was extinguished, and the rat received a pellet in the food receptacle. Then another hole was illuminated, and the same process continued until the rat earned a maximum of 50 pellets in a 30-minutes training. The rats were trained until they reached this training criteria.

#### Training 3

Training 3 was similar to Training 2, except that this time, a delay of 5 seconds was introduced after the beginning of the trial and before one of the holes was illuminated. From Training 3 onwards, correct and incorrect responses were recorded. A correct response was a nosepoke in the illuminated hole and an incorrect response was a nosepoke in any of the other four unlit holes. After each correct or incorrect response, a new trial began with a nosepoke in the food receptacle, a 5 second waiting period and illumination of a random nosepoke hole. Rats had to learn this step until they reached 50 correct responses during the 30minute training.

#### Training 4

For Training 4, after the start of the trial by a nosepoke in the food receptacle, like training 3, a delay of 5 seconds was introduced but now it was additionally signified by an auditory tone. After the tone, one of the five holes was illuminated for a period of 2 seconds. A response during the tone and before receptacle illumination was measured as a premature response but there were no programmatic consequences. Correct and incorrect responses were recorded as well as omission responses, defined as the absence a nosepoke during the allowed 5 seconds after the cue light was illuminated. To advance to the final Training 5, the rats had to make at least 30 correct responses during the 30-minute session. Two sessions were held per day and rats usually required one day to learn this task. However, rats some required a second day of training to achieve the criterion.

#### Training 5

Training 5 (Test phase) was similar to Training 4 except that responses in any of the five holes during the tone were recorded as a premature (as in Training 4), and the rats “punished” with a 5-second time out, where the house light was illuminated. The correct, incorrect and omission responses were recorded as previously mentioned. Training 5 was once per day until reaching criterion in two consecutive sessions. The criteria were as follows: > 20 correct responses, >65% accuracy (correct X 100 /[correct + incorrect]), and <20% omission responses (omissions X 100/ [correct + incorrect + omissions + premature]).

The rats had to reach this criterion two sessions in a row and the second session was considered as the baseline session.

The premature responses were calculated as premature responses X 100 / [correct + incorrect + omissions + premature].

After reaching the criteria in Training 5, the duration of waiting (and the tone) was increased to 7s and then 10s.

### Attention Deficit (AD) task

The procedure was similar to Training 5 of the 5CSRTT, except that there was no tone during the waiting period and hence any response during this period had no programmatic consequences. Therefore, no responses were recorded as premature, even if the rats performed a nose poke in any of the five apertures before the light stimulus appeared in any holes. From session to session the time of illumination was decreased from 3s to 1s, 0.5s and finally 0.2s. The following parameters were evaluated.

% correct responses = correct X 100 / (correct + incorrect + omission)

% incorrect responses = incorrect X 100 / (correct + incorrect + omission)

% omission responses = omission X 100 / (correct + incorrect + omission)

### Delay Discounting task

The delayed discounting task was performed in operant chambers (32 x 24 x 25cm) (Med associates, Inc, St. Albans, VT, USA). Each chamber was individually enclosed in a wooden cubicle equipped with an exhaust fan and a white noise generator. Each operant chamber had a steel grid and a tray with bedding to collect the waste. On the right side of the chamber a food receptacle was present measuring 5 x 5 cm and 2 cm above the grid. Sucrose food pellets of 45mg were used as rewards. One retractable lever was available on either side of the food tray. There were lights in the food receptacle, above each lever, as well as an overhead house light. The rats were trained to lever press for one sucrose pellet (small reward) and to press the opposite lever to obtain four sucrose pellets (large reward). The small reward was always delivered immediately after the lever press. The larger reward was delivered after varying delays following the instrumental response. With this task the rat’s willingness to wait for a larger reward rather than the immediate small reward is measured. In the first training stage rats were trained to lever press for a single pellet, followed by a training period during which the rats were required to wait increasing periods for small rewards. Following this, the delays and large reward were introduced. The second step of the training was similar to the initial training except that a 40 second ITI was introduced irrespective of the animal’s response. The trial started with the house light and stimulus light illuminated to signal a nose poke in the food receptacle. Then, the stimulus light was turned off and either the right or the left lever extended. Once pressed, the lever retracted, and a food pellet dropped in the food receptacle as a reward. In the last training phase, rats learned that one lever was associated with the immediate small reward (1 pellet) the other lever associated with the larger reward (4 pellets), delivered after varying delays. The training was divided into 5 blocks in which each block was composed of 10 trials at the same delay for the larger reward. Each delay was introduced in an increasing order: 0s, 10s, 20s, 40s, and 60s. In between each block, one trial for each lever was run in which choice on this lever was rewarded without a delay. This was done to reinforce the memory of the reward values for each lever between different blocks. Each trial started with the house light and stimulus light ON indicating the appropriate lever and food receptacle for a nose poke. After the nose poke in the food receptacle, both levers extended, and the rat could choose between the left lever which always delivered the small immediate reward (1 pellet) and the right lever which gave the larger reward after varying delays. Upon pressing a lever both levers were retracted and the corresponding reward was delivered. From the nose poke at the beginning of each trial to the next trial the ITI was set to be 100 seconds. If the rats did not perform a nose poke in the 10 seconds following the beginning of the trial or if they did not press any of the two levers in the first 10 seconds after they were extended, the levers were retracted, and the rat did not receive reward. If the rat did not respond, it was counted as an omission. This training session was performed for 16 consecutive days and performance on the final four days was used as a measure of their impulsive choice.

### Saccharin preference test

Rats were single housed in standard laboratory polycarbonate cages (370×260×180mm) where they were tested for saccharin preference. On the first day, animals were given access to one bottle containing 0.2% saccharin solution and one bottle containing water. On the next day the location of the two bottles was exchanged to avoid side preference. Consumption of water and saccharin solutions was recorded daily on each day at the same hour. The following two variables were measured:

Percentage choice for saccharine = Amount of saccharin solution consumed X 100 /(Total amount of liquid consumed)
Saccharin to water ratio = Amount of saccharin consumed / amount of water consumed

### Progressive ratio test

Pressing the right lever once delivered 0.1mL of 0.2% saccharin solution and the rats were trained for 15 days to self-administer the saccharin solution. Following this, the rats were trained on a progressive ratio schedule in which and increasing number of active lever presses were required based on the progression sequence given by the following formula: response ratio = (5e^(reward× 0.2)^) – 560. Hence, the progressive-ratio schedule followed the progression: 1, 2, 4, 6, 9, 12, 15, 20, 25, 32, 40, 50, 62, 77, 95, 118, etc. Each session lasted 90 min, or was stopped following 30 consecutive minutes of inactivity on the active lever. The maximal number of active lever presses was defined as the breakpoint, reflecting the motivation for reward. The breakpoint for each rat was averaged across 3 consecutive daily sessions.

### Compulsivity test

Each lever press delivered 0.1 mL of 0.2% saccharin solution followed by a mild electric foot shock (0.22 mA for 0.5 second) through the grid of the SA chamber when the dipper retracted.

### Goal-directed behavior

Rats in the goal-directed behavioral task were food restricted to 85% of their normal body weight. This experiment was conducted in a self-administration box in which fluid reward could be delivered in a 2-well metallic drinking cup, placed between two operant levers, that allowed for up to 2 solutions to be administered upon the pressing of the appropriate corresponding lever.

#### Training Step 1

Rats were trained on an FR1 schedule where only one lever was available in the self-administration box and one lever press delivered 0.1mL of 0.2% saccharine for a maximum of 50 rewards. This was done alternately for both the levers on separate days for 2 days on each lever.

#### Training Step 2

In this training, lasting 30 mins, every 3 minutes one of two levers was inserted in the chamber and a single lever press (FR1) delivered 0.1 mL of liquid reward. At the end of this 3 min period, the alternate lever was inserted for 3 min, and this cycle repeated 5 times. The reward delivered was either 10% Sucrose or 10% Maltodextrin which was randomized between left and right lever. This step was done for 3 days

#### Training Step 3

This was like step 2, expect that instead of FR1, reward was delivered at random ratio 5 schedule for 3 days

#### Training Step 4

This was like Step 2, expect that instead of FR1, reward was delivered at random ratio 10 schedule for 3 days. The average number of lever presses per min on each lever was calculated and averaged over the last two sessions. If a rat showed excessive side preference by ignoring one lever (<2 lever presses/min) or showed very low baseline lever pressing on either lever (<2 lever presses/min) were excluded from the analysis.

#### Training Step 5

Devaluation: Rats were isolated for 1 hour in a single cage and one of the rewards (sucrose/maltodextrin) was provided ad libitum for a period of 1 hr. The total amount of fluid consumed was measured.

#### Training Step 6

Extinction: Rats were placed in the self-administration boxes and given access to both levers for 5 mins with lever presses having no consequence. Lever pressing behavior was measured as number of lever presses/min.

#### Training Step 7

Consumption test: Rats were reintroduced to the single housed cage and then another bottle of the reward which was not devalued in step 5 was also introduced. Rats had access to 2 bottles for 1 hr and the total fluid consumed was measured.

For experiments using a Latin squares design, rats underwent Training Steps 1-6. They were returned to Training Step 4. Training Step 5 devaluation was done with the reward not used for devaluation previously. This was followed by a return to Training Step 6 and then, Training step 7 (consumption test).

### Electrophysiology experiments

#### Brain Slice Preparation

Animals were anesthetized with isoflurane and decapitated using a guillotine. Brains were extracted and transferred to an ice-cold cutting solution containing (in mM): N-methyl-D-glucamine (NMDG), 93; KCl, 2.5; NaH2PO4, 1.2; NaHCO3, 30; HEPES, 20; Glucose, 25; Ascorbic acid, 5; Sodium pyruvate, 3; MgCl2, 10; CaCl2, 0.5. The brain tissue was cut perpendicular to its longitudinal axis using a razor blade, and then the cut surface glued to the stage of the vibrating tissue slicer (Leica VT1200S, Leica Biosystems). Coronal sections (250 μm) were collected from each brain. The brain slices were then transferred to an oxygenated (95% O2/5% CO2) holding chamber filled with HEPES containing artificial cerebrospinal fluid (aCSF) (in mM: NaCl, 109; KCl, 4.5; NaH2PO4, 1.2; NaHCO3, 35; HEPES, 20; Glucose, 11; Ascorbic acid, 0.4; MgCl2, 1; CaCl2, 2.5) at room temperature (22°C) for at least 30 min.

#### In Vitro Electrophysiology

Hemi-sectioned slices were transferred to a recording chamber (RC-26; Warner Instruments) continuously perfused (2 mL/min) with oxygenated aCSF (in mM: NaCl, 126; KCl, 4.5; NaH2PO4, 1.2; NaHCO3, 26; Glucose, 11; MgCl2, 2; CaCl2, 2.5) using a peristaltic pump (Cole-Parmer). The temperature was maintained at 30–32°C using an inline solution heater connected to a temperature controller (TC-344C/SH-27B, Warner Instruments). Appropriate neurons were identified using an upright videomicroscope (BX51WI, Olympus) equipped with differential interference contrast imaging, and 900 nm infrared illumination. Recording electrodes were fabricated using borosilicate glass (Sutter Instruments, 1.5 mm O.D. × 0.86 mm i.d.) using a horizontal puller (P-97; Sutter Instruments) and filled with a potassium-based internal solution (in mM: K-gluconate, 140; KCl, 5; HEPES, 10; EGTA, 0.2; MgCl2, 2; Mg-ATP, 4; Na2-GTP, 0.3; Na2-phosphocreatine, 10) neutralized to a pH of 7.2 using potassium hydroxide. Electrode resistances were 3–5 MΩ. Whole-cell patch clamp recordings were performed using an Axopatch 200B amplifier (Molecular Devices). Unless otherwise noted, cells were voltage-clamped at −60 mV (with a calculated junction potential of −14 mV). Stimulation protocols and recordings were performed using WinLTP software (WinLTP Ltd, Bristol, UK) and an A/D board (National Instruments, PCI-6251) housed in a personal computer. Hyperpolarizing voltage steps (−10 mV) were delivered via the recording electrode every 30s to monitor whole-cell access and series resistance. Cells with an access change of more than 20% were excluded from analysis.

##### Input resistance

Hyperpolarizing voltage steps (−10 mV, 100ms) were delivered via the recording electrode every 30 s to monitor the steady state input resistance, obtained over the last 20ms of the step. The input resistance was calculated as the average of the last 10 sweeps in voltage clamp.

##### Rheobase

Rheobase was evaluated in current clamp. For the BLA pyramidal neurons and NAc core medium spiny neurons, depolarizing current steps were applied (0–250 pA, 1 sec duration in 10 pA increments, repeated 3 times) from a membrane potential of −60 mV. For PLC pyramidal neurons rheobase was evaluated by applying depolarizing current steps (0–2000 pA, 50 pA increments, 5 ms duration, repeated 3 times) from a membrane potential of −60 mV.

##### Excitability

To measure excitability, depolarizing current steps were applied (0–350 pA (for BLA), (0-250 pA for PLC) and (0-200 pA for NAc core), 50 pA increments, 1s, repeated 3 times from a membrane potential of −60 mV. The mean number of action potentials (APs) during each depolarization step was plotted against the amount of current injected.

##### Afterhyperpolarization (AHP)

The medium and slow AHPs (mAHP and sAHP) were measured in response to a train of 5 depolarizing current pulses (10 ms duration at 700 pA, evoking a train of 5 APs), from a membrane potential of −60mV. Only AHPs elicited by 1 AP per 10ms depolarization pulse were used for analysis. The train was repeated 3 times (10–20s intervals) and the average AHP response was measured. The mAHP (at AHP peak) and the sAHP (270–300ms after the step) amplitudes were analysed separately. The AHP amplitude was measured by subtraction from the preburst baseline membrane potential using custom software (WinWCP v 5.42, courtesy of Dr. John Dempster, Strathclyde University, Glasgow UK).

##### Spontaneous excitatory post synaptic currents (sEPSCs)

The cells were held in voltage clamp at −60mV and continuous recordings were obtained for 1 minute epochs. Detection and analysis of sEPSCs was performed off-line using the MiniAnalysis program (v 6.0.7, Synaptosoft, Inc.), and sEPSCs were visually inspected to confirm events.

### Optogenetic electrophysiology experiments

Virus injections: Rats were anesthetized with isoflurane using a precision vaporizer connected to a nose cone apparatus and injected (1 μl over 10 min) with AAV5-CaMKII-ChR2-eYFP (4.6×1012 virus molecules/ml; University of North Carolina vector core) into Prelimbic cortex (AP: +3.3; ML: ±0.8; DV: −5.0), or BLA (AP: −2.8; ML: ±5.2; DV: −8.0) using a 10-μl Hamilton syringe, an UltraMicroPump, and SYS-Mico4 controller (WPI, Sarasota, FL). Incisions were closed with absorbable sutures, and body temperature was maintained with a heating pad until anesthesia recovery. Postoperative analgesia was done with Meloxicam injection (1mg/kg) subcutaneously for three days.

#### Light-activated EPSCs

Neurons were held at −60mV. Light-activated optical excitatory post-synaptic currents (oEPSCs) were evoked using 473-nm LED pulses (5 ms duration) driven by a Thor Labs (Newton, N.J., USA) LED driver. The LED driver was connected to a 400 μm core, 0.39 NA optical patch cable with its stripped end positioned above the recording location using a micromanipulator. A Thor Labs power meter and sensor was used to determine the laser power measured at the fiber tip across the range of intensity settings (0-1000 mA). In NAc recordings, picrotoxin (100 μM) was used to block GABA_A_ receptor chloride channels. For BLA/PLC recordings, picrotoxin was not used because it caused large, unclamped excitatory synaptic currents that were likely due to disinhibition of recurrent collaterals. To maintain consistency across slices and brain areas, the same injection volume of ChR2 virus (1 μl) and the same duration to permit transfection (6-8 weeks) were used in all experiments. Input-output (I-O) relationships between oEPSC amplitude and laser intensity (2, 3, 4, 5, 6, 7, 8, 9 and 10 mW) were constructed with an average of three oEPSCs generated at each LED intensity. Ratios of pairs of oEPSCs (paired-pulse ratio; PPR) were collected using two laser stimuli with 40, 60, 80, 100, or 200 ms inter-stimulus intervals (ISI), collected at maximum LED intensity (10 mW). The ratio was calculated by averaging 3 consecutive responses and dividing the peak amplitude of the second oEPSC by the first.

Light-activated EPSPs (oEPSPs) and action potentials: oEPSP experiments were done in current clamp with minimal current injected to maintain the cell membrane potential at −60mV. Maximum LED intensity (10 mW) was used to ascertain if the oEPSP generated an action potential.

##### Paired-pulse ratio

Neurons were held at −60mV. oEPSCs were evoked using 473-nm LED pulses (5 ms duration) driven by a Thor Labs LED driver. Two LED pulses at laser intensity 10mW were delivered at an inter-stimulus interval of 40, 60,80,100 and 200 ms. The ratio was calculated by averaging 3 consecutive responses and dividing the peak amplitude of the second oEPSC by the first. This was done three times at each inter-stimulus interval.

##### Spike timing-dependent plasticity

The neuron was held in voltage clamp at −60mV and the average amplitude of oEPSCs at laser intensity 10mW was measured over 5 minutes (every 30 seconds). The neuron was then held in current clamp at −60mV and 100 pulses (1 every second) at laser intensity 10mW followed by a depolarizing current (which was determined by a rheobase experiment) to fire an action potential (1 every second). The neuron was then returned voltage clamp at −60mV and the amplitude of oEPSC measured over 30 mins (every 30 seconds).

### Chemogenetics virus injections

Rats were anesthetized with isoflurane using a precision vaporizer connected to a nose cone apparatus. There were injected intraperitoneally, 3mg/kg of buprenorphine preoperatively and then infused (1μl over 15 min) with AAV8 excitatory DREADD (pAAV-CaMKIIa-hM3D(Gq)-mCherry) (4.6×10^12^ virus molecules/ml; University of North Carolina vector core) in the prelimbic cortex (AP: +3.3; ML: ±0.8; DV: −5.0), using a 5-μl Hamilton syringe, an UltraMicroPump, and SYS-Mico4 controller (WPI, Sarasota, FL). Incisions were closed with absorbable sutures, and body temperature was maintained with a heating pad until anesthesia recovery. Rats were given 3 weeks to recover and tested on the behavioral tasks.

## References

Adriani, W., and Laviola, G. (2003). Elevated levels of impulsivity and reduced place conditioning with d-amphetamine: two behavioral features of adolescence in mice. Behav Neurosci 117, 695–703.

Andersen, S.L., and Teicher, M.H. (2008). Stress, sensitive periods and maturational events in adolescent depression. Trends Neurosci 31, 183–191. 10.1016/j.tins.2008.01.004.

Andersen, S.L., and Teicher, M.H. (2009). Desperately driven and no brakes: developmental stress exposure and subsequent risk for substance abuse. Neurosci Biobehav Rev 33, 516–524. 10.1016/j.neubiorev.2008.09.009.

Andrzejewski, M.E., Schochet, T.L., Feit, E.C., Harris, R., McKee, B.L., and Kelley, A.E. (2011). A comparison of adult and adolescent rat behavior in operant learning, extinction, and behavioral inhibition paradigms. Behav Neurosci 125, 93–105. 10.1037/a0022038.

Arnett, J.J. (1999). Adolescent storm and stress, reconsidered. Am Psychol 54, 317–326.

Arruda-Carvalho, M., Wu, W.C., Cummings, K.A., and Clem, R.L. (2017). Optogenetic Examination of Prefrontal-Amygdala Synaptic Development. J Neurosci 37, 2976–2985. 10.1523/JNEUROSCI.3097-16.2017.

Baarendse, P.J., Counotte, D.S., O’Donnell, P., and Vanderschuren, L.J. (2013). Early social experience is critical for the development of cognitive control and dopamine modulation of prefrontal cortex function. Neuropsychopharmacology 38, 1485–1494. 10.1038/npp.2013.47.

Baker, L.M., Williams, L.M., Korgaonkar, M.S., Cohen, R.A., Heaps, J.M., and Paul, R.H. (2013). Impact of early vs. late childhood early life stress on brain morphometrics. Brain Imaging Behav 7, 196–203. 10.1007/s11682-012-9215-y.

Balleine, B.W., and Dickinson, A. (1998). Goal-directed instrumental action: contingency and incentive learning and their cortical substrates. Neuropharmacology 37, 407–419. 10.1016/s0028-3908(98)00033-1.

Barch, D.M., Belden, A.C., Tillman, R., Whalen, D., and Luby, J.L. (2018). Early Childhood Adverse Experiences, Inferior Frontal Gyrus Connectivity, and the Trajectory of Externalizing Psychopathology. J Am Acad Child Adolesc Psychiatry 57, 183–190. 10.1016/j.jaac.2017.12.011.

Bick, J., and Nelson, C.A. (2016). Early Adverse Experiences and the Developing Brain. Neuropsychopharmacology 41, 177–196. 10.1038/npp.2015.252.

Bourgeois, J.P., Goldman-Rakic, P.S., and Rakic, P. (1994). Synaptogenesis in the prefrontal cortex of rhesus monkeys. Cereb Cortex 4, 78–96. 10.1093/cercor/4.1.78.

Buschman, T.J., and Miller, E.K. (2014). Goal-direction and top-down control. Philos Trans R Soc Lond B Biol Sci 369. 10.1098/rstb.2013.0471.

Caballero, A., Thomases, D.R., Flores-Barrera, E., Cass, D.K., and Tseng, K.Y. (2014). Emergence of GABAergic-dependent regulation of input-specific plasticity in the adult rat prefrontal cortex during adolescence. Psychopharmacology (Berl) 231, 1789–1796. 10.1007/s00213-013-3216-4.

Casey, B.J., Galvan, A., and Somerville, L.H. (2016). Beyond simple models of adolescence to an integrated circuit-based account: A commentary. Dev Cogn Neurosci 17, 128–130. 10.1016/j.dcn.2015.12.006.

Casey, B.J., and Jones, R.M. (2010). Neurobiology of the adolescent brain and behavior: implications for substance use disorders. J Am Acad Child Adolesc Psychiatry 49, 1189–1201; quiz 1285. 10.1016/j.jaac.2010.08.017.

Cauffman, E., Shulman, E.P., Steinberg, L., Claus, E., Banich, M.T., Graham, S., and Woolard, J. (2010). Age differences in affective decision making as indexed by performance on the Iowa Gambling Task. Dev Psychol 46, 193–207. 10.1037/a0016128.

Cunningham, M.G., Bhattacharyya, S., and Benes, F.M. (2002). Amygdalo-cortical sprouting continues into early adulthood: implications for the development of normal and abnormal function during adolescence. J Comp Neurol 453, 116–130. 10.1002/cne.10376.

Doremus-Fitzwater, T.L., and Spear, L.P. (2016). Reward-centricity and attenuated aversions: An adolescent phenotype emerging from studies in laboratory animals. Neurosci Biobehav Rev 70, 121–134. 10.1016/j.neubiorev.2016.08.015.

Eiland, L., Ramroop, J., Hill, M.N., Manley, J., and McEwen, B.S. (2012). Chronic juvenile stress produces corticolimbic dendritic architectural remodeling and modulates emotional behavior in male and female rats. Psychoneuroendocrinology 37, 39–47. 10.1016/j.psyneuen.2011.04.015.

Figner, B., Mackinlay, R.J., Wilkening, F., and Weber, E.U. (2009). Affective and deliberative processes in risky choice: age differences in risk taking in the Columbia Card Task. J Exp Psychol Learn Mem Cogn 35, 709–730. 10.1037/a0014983.

Giedd, J.N. (2004). Structural magnetic resonance imaging of the adolescent brain. Ann N Y Acad Sci 1021, 77–85. 10.1196/annals.1308.009.

Gill, T.M., Castaneda, P.J., and Janak, P.H. (2010). Dissociable roles of the medial prefrontal cortex and nucleus accumbens core in goal-directed actions for differential reward magnitude. Cereb Cortex 20, 2884–2899. 10.1093/cercor/bhq036.

Hare, T.A., Tottenham, N., Galvan, A., Voss, H.U., Glover, G.H., and Casey, B.J. (2008). Biological substrates of emotional reactivity and regulation in adolescence during an emotional go-nogo task. Biol Psychiatry 63, 927–934. 10.1016/j.biopsych.2008.03.015.

Huttenlocher, P.R., de Courten, C., Garey, L.J., and Van der Loos, H. (1982). Synaptogenesis in human visual cortex--evidence for synapse elimination during normal development. Neurosci Lett 33, 247–252. 10.1016/0304-3940(82)90379-2.

Kalivas, P.W., Volkow, N., and Seamans, J. (2005). Unmanageable motivation in addiction: a pathology in prefrontal-accumbens glutamate transmission. Neuron 45, 647–650. 10.1016/j.neuron.2005.02.005.

Kelley, A.E., and Berridge, K.C. (2002). The neuroscience of natural rewards: relevance to addictive drugs. J Neurosci 22, 3306–3311. 20026361.

Kim, P., Evans, G.W., Angstadt, M., Ho, S.S., Sripada, C.S., Swain, J.E., Liberzon, I., and Phan, K.L. (2013). Effects of childhood poverty and chronic stress on emotion regulatory brain function in adulthood. Proc Natl Acad Sci U S A 110, 18442–18447. 10.1073/pnas.1308240110.

Leussis, M.P., Lawson, K., Stone, K., and Andersen, S.L. (2008). The enduring effects of an adolescent social stressor on synaptic density, part II: Poststress reversal of synaptic loss in the cortex by adinazolam and MK-801. Synapse 62, 185–192. 10.1002/syn.20483.

McLaughlin, K.A., Peverill, M., Gold, A.L., Alves, S., and Sheridan, M.A. (2015). Child Maltreatment and Neural Systems Underlying Emotion Regulation. J Am Acad Child Adolesc Psychiatry 54, 753–762. 10.1016/j.jaac.2015.06.010.

Nooner, K.B., Mennes, M., Brown, S., Castellanos, F.X., Leventhal, B., Milham, M.P., and Colcombe, S.J. (2013). Relationship of trauma symptoms to amygdala-based functional brain changes in adolescents. J Trauma Stress 26, 784–787. 10.1002/jts.21873.

Novick, A.M., Miiller, L.C., Forster, G.L., and Watt, M.J. (2013). Adolescent social defeat decreases spatial working memory performance in adulthood. Behav Brain Funct 9, 39. 10.1186/1744-9081-9-39.

Padmanabhan, V., Cardoso, R.C., and Puttabyatappa, M. (2016). Developmental Programming, a Pathway to Disease. Endocrinology 157, 1328–1340. 10.1210/en.2016-1003.

Petanjek, Z., Judas, M., Kostovic, I., and Uylings, H.B. (2008). Lifespan alterations of basal dendritic trees of pyramidal neurons in the human prefrontal cortex: a layer-specific pattern. Cereb Cortex 18, 915–929. 10.1093/cercor/bhm124.

Phillipson, O.T., and Griffiths, A.C. (1985). The topographic order of inputs to nucleus accumbens in the rat. Neuroscience 16, 275–296. 10.1016/0306-4522(85)90002-8.

Rich, E.L., and Romero, L.M. (2005). Exposure to chronic stress downregulates corticosterone responses to acute stressors. Am J Physiol Regul Integr Comp Physiol 288, R1628–1636. 10.1152/ajpregu.00484.2004.

Romer, D., Reyna, V.F., and Satterthwaite, T.D. (2017). Beyond stereotypes of adolescent risk taking: Placing the adolescent brain in developmental context. Dev Cogn Neurosci 27, 19–34. 10.1016/j.dcn.2017.07.007.

Schalinski, I., Teicher, M.H., Carolus, A.M., and Rockstroh, B. (2018). Defining the impact of childhood adversities on cognitive deficits in psychosis: An exploratory analysis. Schizophr Res 192, 351–356. 10.1016/j.schres.2017.05.014.

Shao, F., Jin, J., Meng, Q., Liu, M., Xie, X., Lin, W., and Wang, W. (2009). Pubertal isolation alters latent inhibition and DA in nucleus accumbens of adult rats. Physiol Behav 98, 251–257. 10.1016/j.physbeh.2009.05.021.

Smith, S.M., and Vale, W.W. (2006). The role of the hypothalamic-pituitary-adrenal axis in neuroendocrine responses to stress. Dialogues Clin Neurosci 8, 383–395.

Sokolowski, K., and Corbin, J.G. (2012). Wired for behaviors: from development to function of innate limbic system circuitry. Front Mol Neurosci 5, 55. 10.3389/fnmol.2012.00055.

Sotres-Bayon, F., and Quirk, G.J. (2010). Prefrontal control of fear: more than just extinction. Curr Opin Neurobiol 20, 231–235. 10.1016/j.conb.2010.02.005.

Spear, L.P. (2000). The adolescent brain and age-related behavioral manifestations. Neurosci Biobehav Rev 24, 417–463.

Stuber, G.D., Sparta, D.R., Stamatakis, A.M., van Leeuwen, W.A., Hardjoprajitno, J.E., Cho, S., Tye, K.M., Kempadoo, K.A., Zhang, F., Deisseroth, K., and Bonci, A. (2011). Excitatory transmission from the amygdala to nucleus accumbens facilitates reward seeking. Nature 475, 377–380. 10.1038/nature10194.

Telzer, E.H., Ichien, N.T., and Qu, Y. (2015). Mothers know best: redirecting adolescent reward sensitivity toward safe behavior during risk taking. Soc Cogn Affect Neurosci 10, 1383–1391. 10.1093/scan/nsv026.

Thomason, M.E., Marusak, H.A., Tocco, M.A., Vila, A.M., McGarragle, O., and Rosenberg, D.R. (2015). Altered amygdala connectivity in urban youth exposed to trauma. Soc Cogn Affect Neurosci 10, 1460–1468. 10.1093/scan/nsv030.

Torregrossa, M.M., Xie, M., and Taylor, J.R. (2012). Chronic corticosterone exposure during adolescence reduces impulsive action but increases impulsive choice and sensitivity to yohimbine in male Sprague-Dawley rats. Neuropsychopharmacology 37, 1656–1670. 10.1038/npp.2012.11.

Tottenham, N., and Gabard-Durnam, L.J. (2017). The developing amygdala: a student of the world and a teacher of the cortex. Curr Opin Psychol 17, 55–60. 10.1016/j.copsyc.2017.06.012.

Vaghi, M.M., Moutoussis, M., Vasa, F., Kievit, R.A., Hauser, T.U., Vertes, P.E., Shahar, N., Romero-Garcia, R., Kitzbichler, M.G., Bullmore, E.T., et al. (2020). Compulsivity is linked to reduced adolescent development of goal-directed control and frontostriatal functional connectivity. Proc Natl Acad Sci U S A 117, 25911–25922. 10.1073/pnas.1922273117.

Walker, D.M., Bell, M.R., Flores, C., Gulley, J.M., Willing, J., and Paul, M.J. (2017). Adolescence and Reward: Making Sense of Neural and Behavioral Changes Amid the Chaos. J Neurosci 37, 10855–10866. 10.1523/JNEUROSCI.1834-17.2017.

Wassum, K.M., and Izquierdo, A. (2015). The basolateral amygdala in reward learning and addiction. Neurosci Biobehav Rev 57, 271–283. 10.1016/j.neubiorev.2015.08.017.

Watt, M.J., Burke, A.R., Renner, K.J., and Forster, G.L. (2009). Adolescent male rats exposed to social defeat exhibit altered anxiety behavior and limbic monoamines as adults. Behav Neurosci 123, 564–576. 10.1037/a0015752.

Wei, J., Zhong, P., Qin, L., Tan, T., and Yan, Z. (2018). Chemicogenetic Restoration of the Prefrontal Cortex to Amygdala Pathway Ameliorates Stress-Induced Deficits. Cereb Cortex 28, 1980–1990. 10.1093/cercor/bhx104.

Yuan, P., and Raz, N. (2014). Prefrontal cortex and executive functions in healthy adults: a meta-analysis of structural neuroimaging studies. Neurosci Biobehav Rev 42, 180–192. 10.1016/j.neubiorev.2014.02.005.

